# A role for cytoglobin in regulating intracellular hydrogen peroxide and redox signals in the vasculature

**DOI:** 10.1101/2023.03.31.535146

**Authors:** Clinton Mathai, Frances Jourd’heuil, Le Gia Cat Pham, Kurrim Gilliard, Joseph Balnis, Annie Jen, Katherine A. Overmyer, Joshua J Coon, Ariel Jaitovich, Benoit Boivin, David Jourd’heuil

## Abstract

The oxidant hydrogen peroxide serves as a signaling molecule that alters many aspects of cardiovascular functions. Recent studies suggest that cytoglobin – a hemoglobin expressed in the vasculature – may promote electron transfer reactions with proposed functions in hydrogen peroxide decomposition. Here, we determined the extent to which cytoglobin regulates intracellular hydrogen peroxide and established mechanisms. We found that cytoglobin decreased the hyperoxidation of peroxiredoxins and maintained the activity of peroxiredoxin 2 following challenge with exogenous hydrogen peroxide. Cytoglobin promoted a reduced intracellular environment and facilitated the reduction of the thiol-based hydrogen peroxide sensor Hyper7 after bolus addition of hydrogen peroxide. Cytoglobin also limited the inhibitory effect of hydrogen peroxide on glycolysis and reversed the oxidative inactivation of the glycolytic enzyme GAPDH. Our results indicate that cytoglobin in cells exists primarily as oxyferrous cytoglobin (CygbFe^2+^-O_2_) with its cysteine residues in the reduced form. We found that the specific substitution of one of two cysteine residues on cytoglobin (C83A) inhibited the reductive activity of cytoglobin on Hyper7 and GAPDH. Carotid arteries from cytoglobin knockout mice were more sensitive to glycolytic inhibition by hydrogen peroxide than arteries from wildtype mice. Together, these results support a role for cytoglobin in regulating intracellular redox signals associated with hydrogen peroxide through oxidation of its cysteine residues, independent of hydrogen peroxide reaction at its heme center.

## 1. Introduction

The biological functions of mammalian globins such as hemoglobin and myoglobin are to transport and store molecular oxygen and regulate nitric oxide^1^. Biochemical studies over the past 20 years indicate that cytoglobin, a mammalian globin with broad tissue expression, may also be uniquely positioned to support the regulation of several reactive oxygen species^2–4^. For example, cytoglobin reacts with superoxide at a rate that is competitive with conventional superoxide dismutases^5^. Like myoglobin and hemoglobin, cytoglobin also reacts with hydrogen peroxide to form ferryl intermediates suggesting a role for cytoglobin to scavenge hydrogen peroxide^6–8^. However, given that this reaction occurs at a rate that is much slower than known peroxidases such as catalase and heme myeloperoxidase, the role of cytoglobin as a physiologically relevant scavenger of hydrogen peroxide is unclear^9,10^. Past work performed on cultured cell models or *in vivo* depicts a similarly complex role for cytoglobin. Overall, increased cytoglobin expression during oxidative stress^11–15^, and in some cases the reciprocal cytotoxic effect associated with loss of cytoglobin have led to the concept that cytoglobin may prevent oxidative damage from hydrogen peroxide^16–18^. However, the cause of increased oxidative stress following cytoglobin deletion, whether direct – through loss of antioxidant activity - or indirect - via a dysregulated inflammatory response or gene expression profile^17–19^-remains to be clarified.

Given the inconsistencies regarding the reactivity of cytoglobin toward hydrogen peroxide and the ambiguous physiological function of cytoglobin, we designed a series of experiments to clarify whether cytoglobin regulates intracellular hydrogen peroxide in cells and *in vivo*. To this end, we probed three different aspects of the cellular response to hydrogen peroxide. We found that (i) ectopic expression of cytoglobin maintains active peroxiredoxin 2; (ii) it is associated with a more reduced environment and accelerates the rate of reduction of the hydrogen peroxide sensor Hyper7 following hydrogen peroxide treatment; (iii) cytoglobin antagonizes the inhibitory effect of hydrogen peroxide on glycolysis and maintains GAPDH activity. We found that cytoglobin exists intracellularly in the oxyferrous form with its two surface cysteine residues reduced. Furthermore, in mouse carotid arteries, deletion of cytoglobin was associated with a decrease in glycolytic activity following hydrogen peroxide challenge. These results support the hypothesis that cytoglobin can function as an antioxidant by removing hydrogen peroxide primarily through reactions at cysteine 83. These redox properties might account for some of the proposed roles of cytoglobin against oxidative stress.

## 2. Results

### 2.1 Cytoglobin inhibits peroxiredoxin hyperoxidation and maintains active peroxiredoxin 2

A sensitive method to measure changes in intracellular redox state in response to hydrogen peroxide is by examining the conversion of cytosolic peroxiredoxins to different oxidation states^20^. In short, the catalytic cycle of conventional peroxiredoxins consists of the oxidation of a peroxidatic cysteine residue by hydrogen peroxide to yield the initial sulfenic acid intermediate^21^. This promotes peroxiredoxin homodimerization through formation of intermolecular disulfides with a resolving cysteine residue. The disulfide is then reduced by the thioredoxin/thioredoxin reductase system to regenerate the thiolate on the peroxidatic cysteine residue. The catalytic activity of mammalian peroxiredoxins is also accompanied by the progressive oxidation of cysteine sulfenic intermediates to sulfinic and sulfonic acid moieties (termed hyperoxidation), which leads to inactivation of their peroxidase activity^22–24^. We reasoned that if cytoglobin were to promote the degradation of intracellular hydrogen peroxide, this should be competitive with peroxiredoxins.

For these experiments, HEK293 cells were transfected with plasmids to produce human cytoglobin (hCYGB) or a control plasmid (EV, Empty Vector) and selected for stable expression (**Figure 1A**). Hyperoxidation of peroxiredoxins can be monitored on Western blots using an antibody directed against stable sulfinic and sulfonic residues^25^. We found that cytoglobin expression was associated with a decrease in hyperoxidation following treatment of the cells with hydrogen peroxide (**Figure 1A**). Next, we followed the monomeric/dimeric states of the two primary cytosolic peroxiredoxins, peroxiredoxin 1 and 2. This can be achieved through post-treatment of cells with N-ethylmaleimide (NEM) followed by non-reducing SDS PAGE of the lysates. The monomeric and dimeric forms of the two peroxiredoxins were observed in both control and cytoglobin expressing HEK293 cells (**Figure 1B and 1C**). Peroxiredoxin 1 responded sharply to the addition of hydrogen peroxide with an increase in the dimeric form at 2 min (**Figure 1B**). In contrast, dimeric peroxiredoxin 2 was present almost in similar amounts to the monomeric form, in the absence of hydrogen peroxide (**Figure 1C**). Consistent with past studies^26^, active dimers in control cells were progressively lost following the addition of hydrogen peroxide due to progressive hyperoxidation of the monomeric peroxiredoxin. Peroxiredoxin 1 activity was unaffected by cytoglobin expression (**Figure 1B**). However, we found that the loss of dimeric peroxiredoxin 2 was delayed in the presence of cytoglobin with the 2 min time-point reaching statistical significance (**Figure 1C**). Overall, our results indicate that the antioxidant effect of cytoglobin is sensitive enough to alter specifically peroxiredoxin 2 activity through a decrease in the rate of reduction of the active dimeric form.

**Fig. 1.**
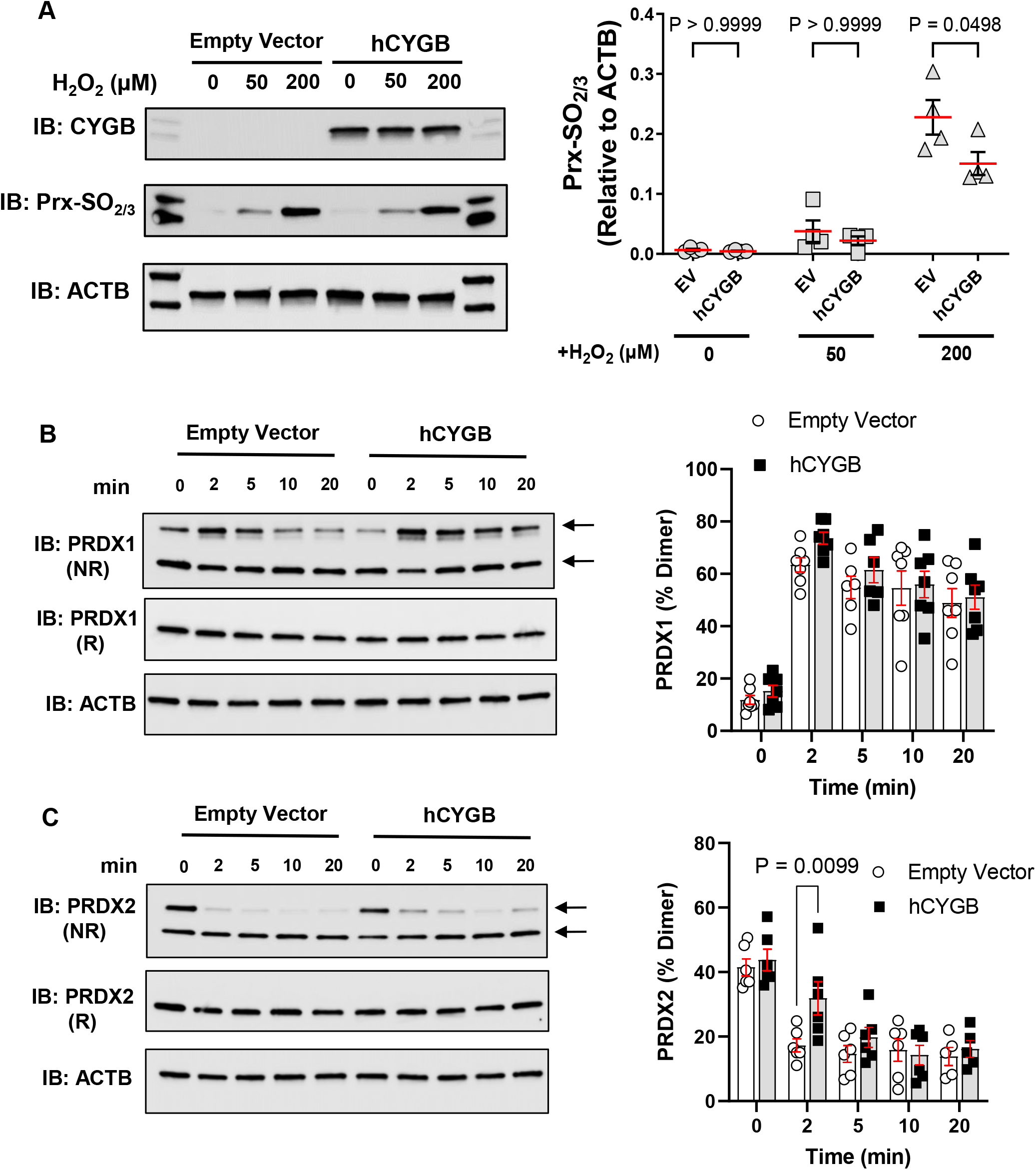
Cytoglobin inhibits peroxiredoxin hyperoxidation and maintains active peroxiredoxin 2. **A,** Control (Empty Vector) and cytoglobin (hCYGB) expressing HEK293 cells were exposed to hydrogen peroxide for 10 min followed by the addition of NEM. Lysates were used for non-reducing SDS PAGE and Western blotting to examine protein levels of hyperoxidized peroxiredoxins (Prx-SO2/3) using beta actin (ACTB) as an internal reference. Cytoglobin levels (CYGB) are also shown. Right panel, quantitation of experiments described in left panel. **B** and **C**, Control (Empty Vector) and cytoglobin (hCYGB) expressing HEK293 cells were exposed to 150 µM hydrogen peroxide for various amounts of time followed by the addition of NEM. Lysates were used in non-reducing SDS PAGE and Western blotting to examine protein levels of peroxiredoxin 1 (B, PRXD1) and 2 (C, PRXD2). Arrows indicate the position of the dimers and monomers. Right panels, quantitation of experiments represented in left panels. All graphs show mean±SEM and each point represents one independent experimental replicate. Statistical analysis was performed using two-way ANOVA.

### 2.2 Cytoglobin promotes the reduction of the hydrogen peroxide sensor Hyper7

In the next set of experiments, we monitored in real time the effect of cytoglobin on intracellular hydrogen peroxide with Hyper7, a genetically encoded hydrogen peroxide sensor (**Figure 2A**). Hyper7 consists of a circularly permutated yellow fluorescent protein (cYFP) that frames the hydrogen peroxide sensing domain of *Neisseria meningitidis* OxyR^27^. This sensing domain contains two cysteine residues that undergo reversible oxidation by hydrogen peroxide to form an intramolecular disulfide bridge. This leads to a change in Hyper7 excitation maxima at 500 and 400 nm that can be quantified and used to calculate ratiometric values that reflect the levels of oxidation of the probe. To minimize the confounding effect of possible topological changes in hydrogen peroxide across different subcellular compartments, we expressed Hyper7-NES (NES, nuclear exclusion sequence) for cytosolic localization^27,28^. We found that the ratiometric baseline values for Hyper7-NES were lower in cytoglobin expressing cells compared to controls suggesting a more reduced intracellular environment with cytoglobin (**Figure 2B and C**). There was a rapid increase in Hyper7-NES oxidation following the addition of hydrogen peroxide in control and cytoglobin expressing cells. Significantly, cytoglobin expression was associated with a sharp increase in Hyper7-NES reduction rates compared to control such that approximately 20 min following the addition of hydrogen peroxide, ratiometric values were returned to baseline (**Figure 2B**). This was most evident at the 100 and 200 µM final concentrations of hydrogen peroxide tested (**Figure 2D**), with no statistically significant effect at 20 and 50 µM.

**Fig. 2.**
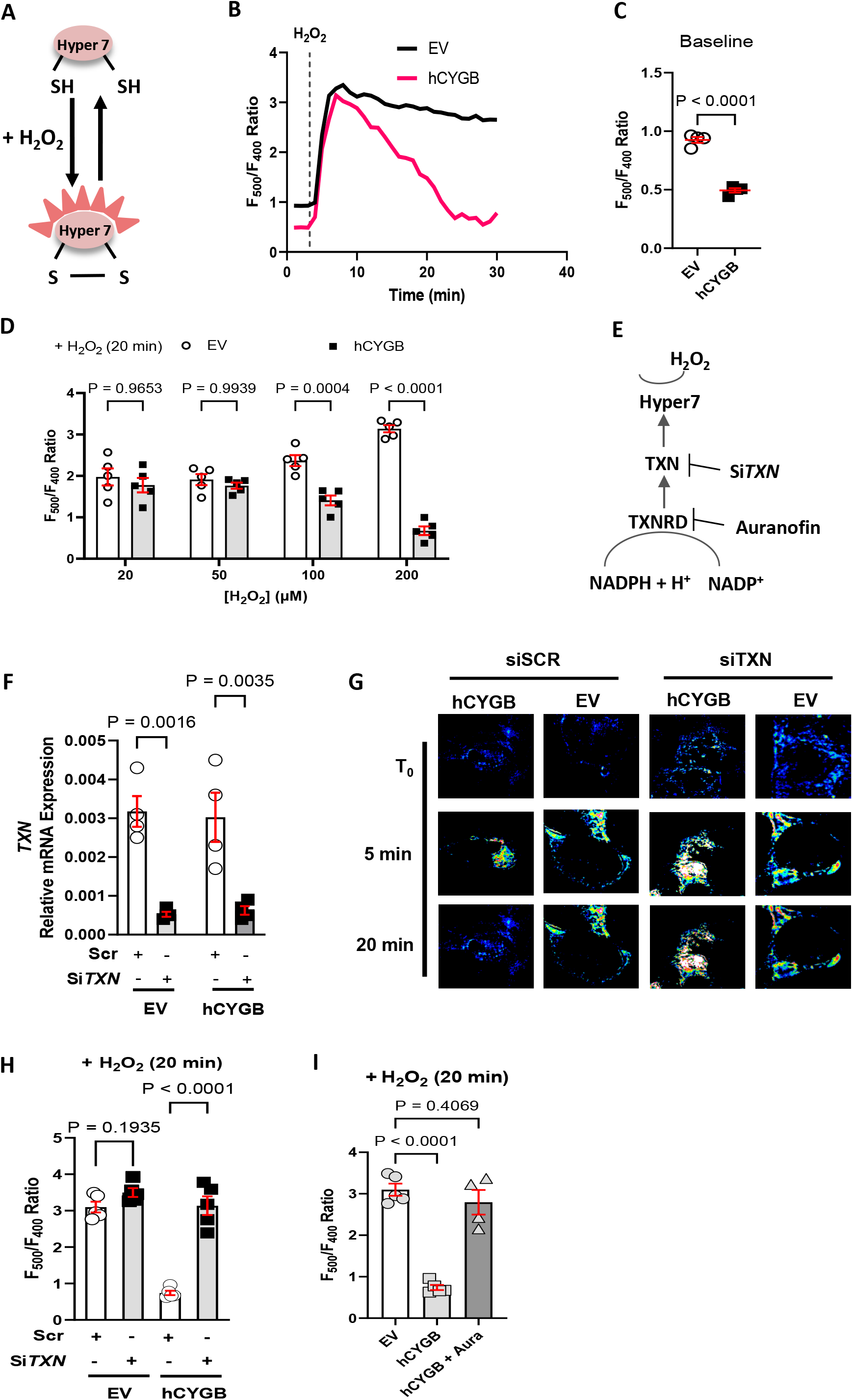
Cytoglobin decreases baseline oxidation of the hydrogen peroxide sensor Hyper7 and facilitates its reduction. **A**, Schematic of Hyper7. The genetically encoded hydrogen peroxide sensor Hyper7 contains two cysteine residues that are reversibly oxidized by hydrogen peroxide, leading to ratiometric changes in fluorescence excitation. **B**, Representative profile of the ratiometric measurement of Hyper7 fluorescence over time in control (EV= Empty Vector) HEK293 cells or cells stably expressing human cytoglobin (hCYGB), both with transient expression of Hyper7-NES (NES, nuclear exclusion signal). Fluorescence intensities were recorded at excitation wavelengths 500 and 400 nm. Following baseline determination, cells were treated with bolus hydrogen peroxide (H_2_O_2_). Profiles are averaged over 4-6 cells. **C**, Quantitation of baseline values for Hyper7 ratiometric fluorescence measurements in control (Empty Vector) and cytoglobin (hCYGB) expressing cells. Each point represents an independent experimental replicate with bar as mean+/− SEM. Statistical analysis was performed using unpaired Student’s t-test. **D**, HEK293 cells were treated with different concentrations of hydrogen peroxide and Hyper7 ratiometric changes were recorded over time and quantified at the 20 min time-point. Each point represent an independent experimental replicate with bar as mean+/− SEM with two-way ANOVA. **E**, Schematic representation of Hyper7 oxidation by hydrogen peroxide and its reduction by the thioredoxin/thioredoxin reductase system. **F**, Control (EV = Empty Vector) HEK293 cells or cells expressing human cytoglobin (hCYGB) were transfected with scrambled (Scr) siRNA or siRNA targeting thioredoxin 1 mRNA (si*TXN*) and relative mRNA expression of *TXN* was determined by quantitative PCR. Each point represents an independent experimental replicate with bar as mean+/− SEM with two-way ANOVA. **G**, Representative pseudo-color images for Hyper7 ratiometric determination Control (EV = Empty Vector) HEK293 cells or cells expressing human cytoglobin (hCYGB) were transfected with scrambled (Scr) siRNA or siRNA targeting thioredoxin 1 mRNA (si*TXN*) followed by transfection of Hyper 7 and treatment with 200 µM hydrogen peroxide. Images shown before addition of hydrogen peroxide (T_0_), and 5 and 20 min after addition. **H**, Quantitation of experiments described in G at the 20 min time-point. Each point represents an independent experimental replicate with bar as mean+/− SEM with two-way ANOVA. **I**, Control (Empty Vector) cells or cells expressing human cytoglobin (hCYGB) were treated with the thioredoxin reductase inhibitor auranofin (Aura) followed by the addition of 200 µM hydrogen peroxide and Hyper 7 ratiometric changes were recorded over time and quantified at the 20 min time-point. Each point represents an independent experimental replicate with bar as mean+/− SEM with one-way ANOVA.

Dynamic changes in Hyper7 fluorescence depend on the rate of hydrogen peroxide decomposition by heme and thiol-based peroxidases but also the reduction rate of Hyper7 by intracellular reducing systems. Past studies indicated that Hyper7 is reduced by the thioredoxin /thioredoxin reductase system^29^ (**Figure 2E**). In agreement, decrease in thioredoxin 1 level (gene code TXN, **Figure 2F**)) through siRNA was sufficient to inhibit cytoglobin-mediated reduction of Hyper7-NES (**Figures 2G and H**). The increase in Hyper7 reduction rate by cytoglobin was also inhibited by pretreatment with the thioredoxin reductase inhibitor auranofin (**Figure 2I**). These results indicated that the reductive effect of cytoglobin relied on a functional thioredoxin/thioredoxin reductase system.

*In vitro*, cytoglobin ligand binding and reactivity require a functional heme center that is stabilized through a proximal and distal histidine residue^30^. In addition, heme chemistry is allosterically regulated by an intramolecular disulfide bridge between Cys38 and Cys83^9^. To determine the role of these structural features, we substituted each of these residues with alanine and generated HEK293 cells stably expressing these variants (**Figure 3A**). Consistent with a role for the heme center, hyper7-NES kinetics following the expression of the H81A (distal histidine residue) or H113A (proximal histidine residue) mutants in HEK cells were not different from the control cells lacking cytoglobin (**Figure 3B**). Most importantly, while the C38A mutant was functional, the C83A mutant also lacked reductive activity towards Hyper7-NES (**Figure 3B**).

**Fig. 3.**
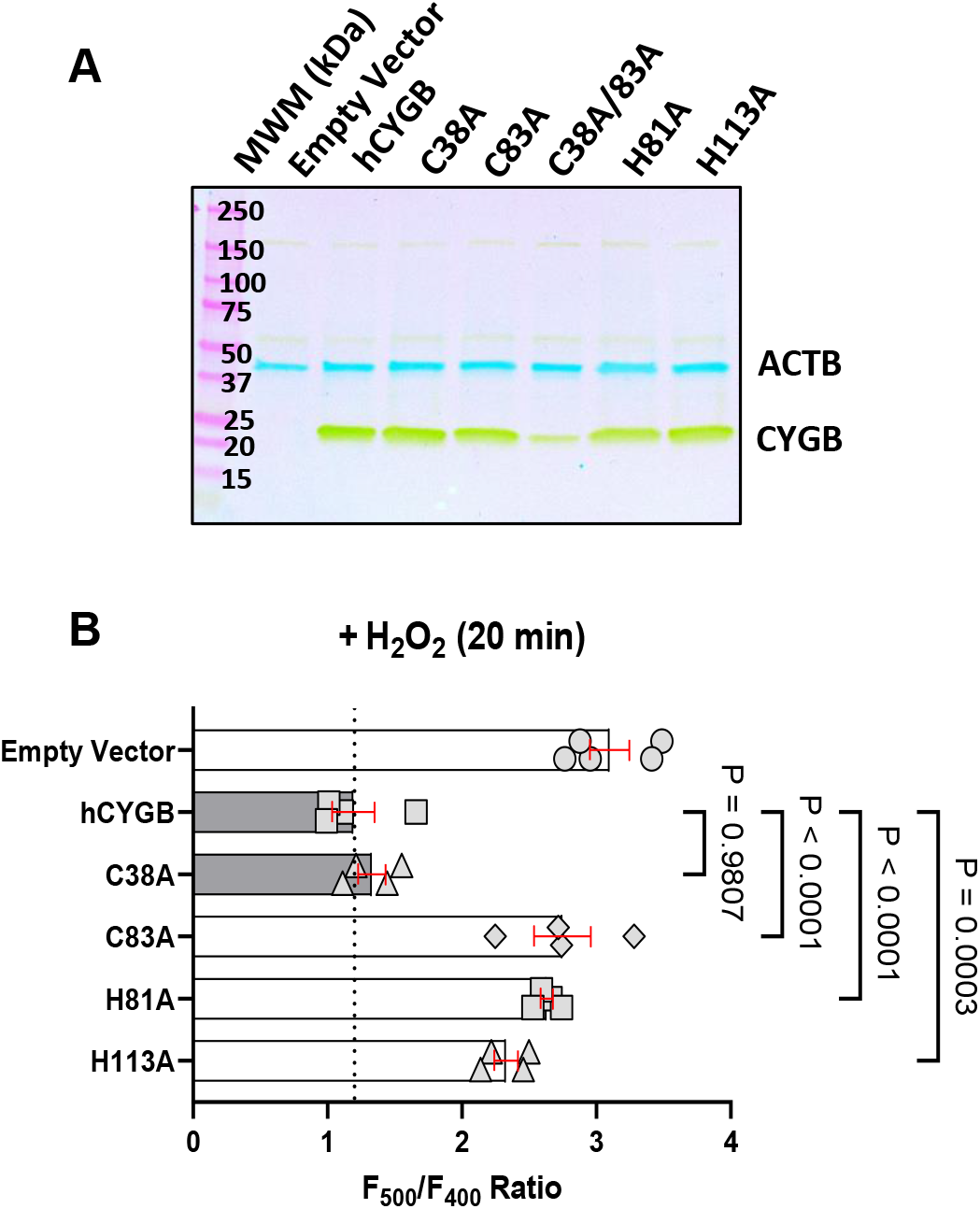
Cytoglobin cysteine 83 promotes Hyper7 reduction. **A**, Western blot analysis of lysates from HEK293 cells stably expressing cytoglobin cysteine and histidine variants using β-actin (gene code ACTB) as an internal reference. **B**, HEK293 cells expressing the cytoglobin variants were transiently transfected with Hyper7-NES followed by treatment with 200 uM hydrogen peroxide. Ratiometric changes were recorded over time and quantified at the 20 min time-point. Mean±SEM with n=4, one-way ANOVA.

### 2.3 Cytoglobin limits the sensitivity of glycolysis to hydrogen peroxide in cultured cells

In addition to peroxiredoxins, past studies have shown that glycolytic enzymes are sensitive to hydrogen peroxide mediated inhibition^31^. We reasoned that examination of glycolytic activity may also serve as a surrogate to test for hydrogen peroxide sensitivity. Since we hypothesized that cytoglobin limits intracellular hydrogen peroxide, we would predict that loss of cytoglobin would result in sensitization of glycolysis to hydrogen peroxide-mediated inhibition (**Figure 4A**). Cellular glycolytic turnover can be assessed by measuring extracellular acidification rates (ECAR) using Seahorse XF technology. We found that addition of bolus hydrogen peroxide inhibited ECAR from HEK293 cells stably transfected with the empty vector in a dose-dependent manner (**Figure 4B**) and this was inhibited upon expression of human cytoglobin, consistent again with a protective effect associated with cytoglobin expression (**Figure 4C and D**). To confirm these results, we performed a targeted metabolomic analysis of glycolytic intermediates in these cells following exposure to bolus hydrogen peroxide for 10 min. Hydrogen peroxide treatment increased all metabolites tested including glucose-6-phosphate, fructose-6-phosphate, fructose-biphosphate, and glyceraldehyde 3 phosphate concentrations (**Figure 4E**). There was also a 5 to 7-fold reduction in the concentration of 2/3 phosphoglycerate compared to glyceraldehyde -3-phosphate, presumably through inhibition of GAPDH. The largest change in metabolites (fructose-biphosphate, and glyceraldehyde 3 phosphate) were statistically significantly decreased in hCYGB expressing compared to control empty vector cells (**Figure 4E**). Hydrogen peroxide treatment also increased levels of ribose-5-phosphate and sedoheptulose-7-phosphate, two intermediates of the pentose monophosphate pathway (**Supplementary Figure 1**). This increase was significantly attenuated upon expression of cytoglobin. Finally, we directly established that GAPDH activity was inhibited following treatment with hydrogen peroxide and cytoglobin expression reversed this effect (**Figure 5**). Importantly, the C83A substitution reversed the effect of cytoglobin while cytoglobin with the C38A substitution was still functional (**Figure 5**). Overall, these results showed an attenuation of the effect of hydrogen peroxide on glycolysis and the pentose monophosphate pathway by cytoglobin. Like its reductive activity measured through Hyper7, the antioxidant effect of cytoglobin was dependent on cysteine 83.

**Fig. 4.**
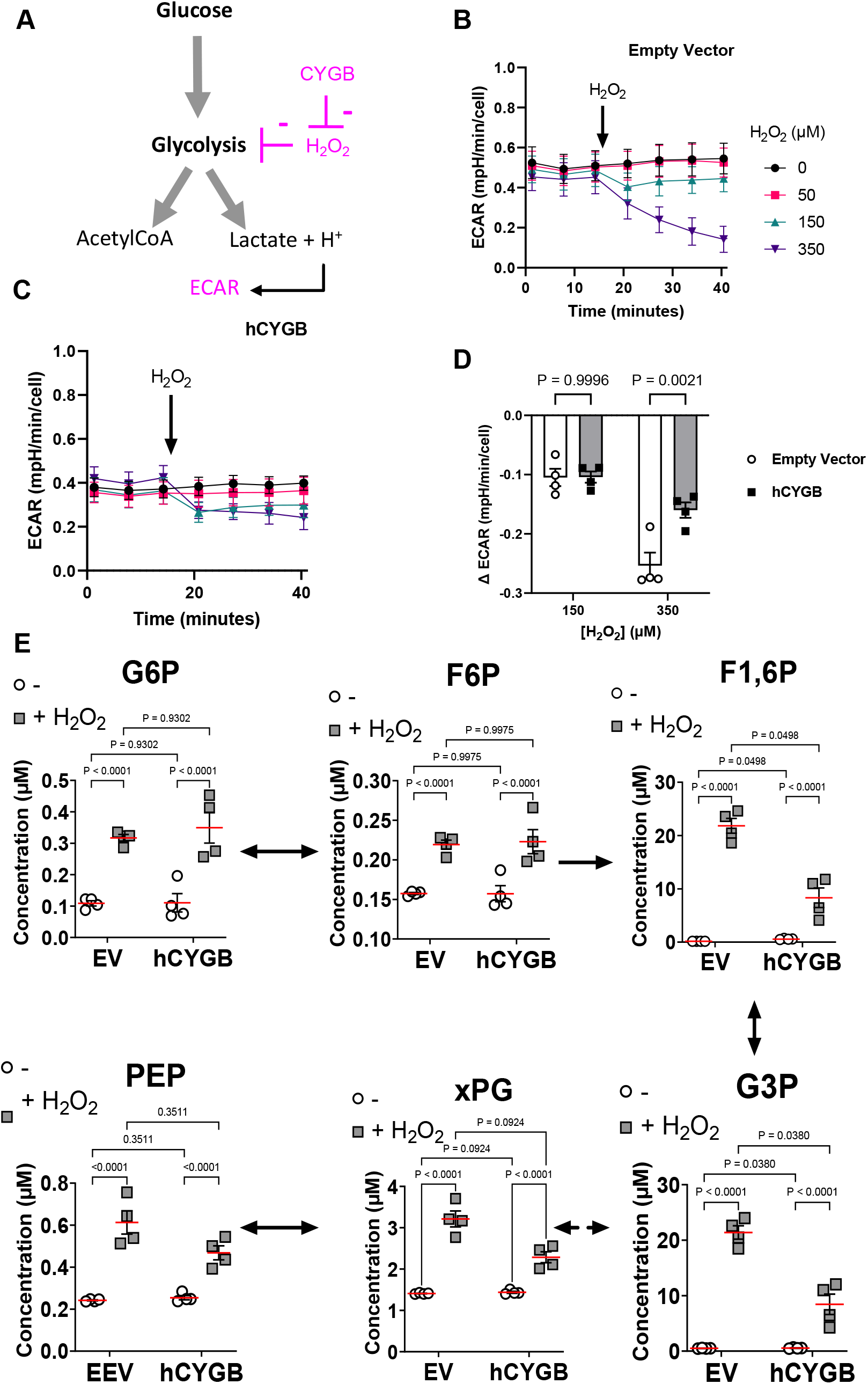
Cytoglobin desensitizes glycolysis from hydrogen peroxide-mediated inhibition in cultured cells. **A**, Schematic of the proposed effect of hydrogen peroxide and cytoglobin on the glycolytic pathway. Following exposure to hydrogen peroxide, glycolysis is inhibited leading to a decrease in extracellular acidification rates (ECAR); if cytoglobin effectively scavenges hydrogen peroxide, we would expect a decrease sensitivity of ECAR following expression of cytoglobin. **B** and **C**, Representative tracing of extracellular acidification rate (ECAR) measurements obtained from HEK293 cells without (B, Empty Vector) or with (C, hCYGB) expression of cytoglobin. Arrow indicates addition of hydrogen peroxide (H_2_O_2_) at different final concentrations. Each time-point and bar represent the average and standard error for at least 4 wells per time point (technical replicates). **D**, Quantitation of the change in ECAR over time following the addition of hydrogen peroxide (H_2_O_2_, 150 and 350 μM). Each point represents independent experimental replicates. Statistical analysis was performed using two-way ANOVA. **E**, Metabolic response of control HEK293 cells (EV = Empty Vector) and HEK293 cells expressing human cytoglobin (hCYGB) following exposure to 150 µM hydrogen peroxide for 10 minutes. Concentrations of intracellular glycolytic metabolites was determined by mass spectrometry. Abbreviations: G6P, glucose-6-phosphate; F6P, fructose-6-phosphate; F1, 6P, Fructose-1-6-biphosphate; G3p, Glucose-3-phosphate; xPG, 2/3-phosphoglycerate; PEP, phosphoenolpyruvate. All graphs show mean±SEM of four independent experimental replicates. Statistical analysis was performed using two-way ANOVA.

**Fig. 5.**
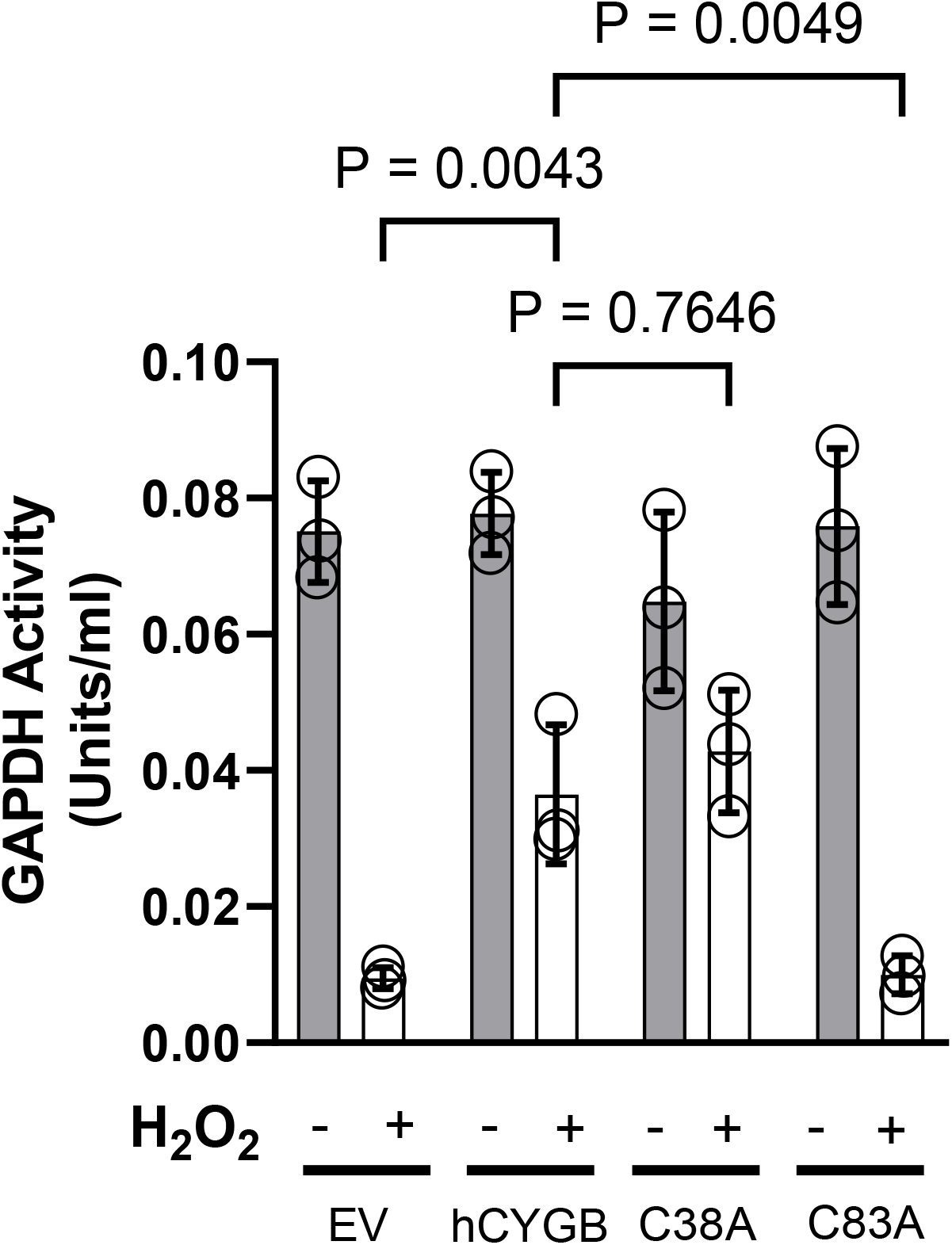
Cytoglobin at Cys83 promotes GAPDH activity in response to hydrogen peroxide. Control (EV = Empty Vector) and cytoglobin (hCYGB) expressing HEK293 cells were exposed to 150 µM hydrogen peroxide for 10 min and lysates were prepared for determination of GAPDH activity. Each point represents independent experimental replicates. Statistical analysis was performed using two-way ANOVA.

### 2.5 The intracellular redox state of cytoglobin

Both cysteine and heme oxidation could contribute to hydrogen peroxide decomposition by cytoglobin. To determine the heme oxidation state of intracellular cytoglobin, we generated spectra on live HEK293 cell suspensions using a spectrophotometer equipped with an integrating cavity. The integrating cavity eliminates scattering losses due to the turbidity of cell suspensions and increases the pathlength with improved detectability. Cells expressing cytoglobin showed characteristic spectral features at the Soret band with shift of the 417m peak to 428 nm following the addition of sodium dithionite (**Figure 6A**). The differential spectra between empty vectors and human cytoglobin expressing cells displayed a distinct Soret peak at 417 nm and primary peaks in the α/β regions at 540 and 578 nm, consistent with oxycytoglobin (CygbFe^2+^-O_2_). Reduction of the cell suspension with dithionite led to the expected shift at 532 and 560 nm upon reduction to deoxycytoglobin (CygbFe^2+^, **Figure 6B**). Next, experiments were performed to examine the reaction of intracellular oxycytoglobin with exogenous hydrogen peroxide. We tested four different concentrations of hydrogen peroxide between 125 µM and 500 µM at cell concentrations ranging from 1×10^6^ to 1×10^7^ cells/ml. Following 10 minutes of hydrogen peroxide treatment no change in the absorption spectra of intracellular oxycytoglobin was observed (**Figure 6C**).

**Fig. 6.**
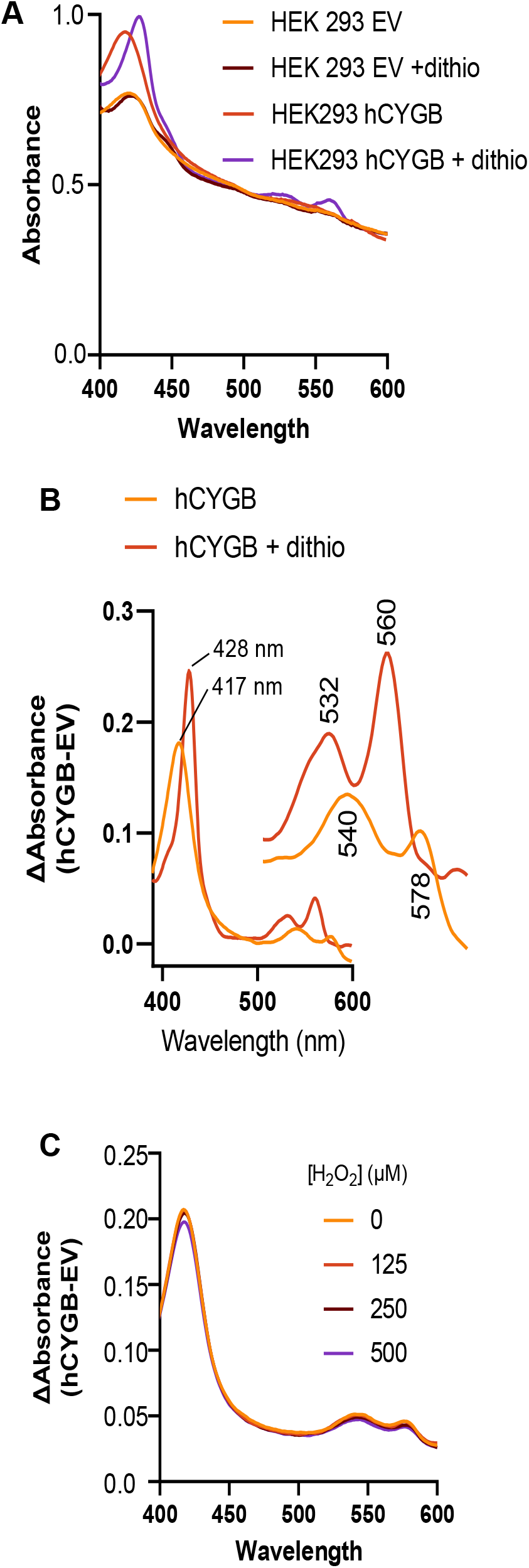
Intracellular oxycytoglobin is unchanged by hydrogen peroxide. **A,** Spectra from HEK293 cells expressing human cytoglobin (hCYGB) and with empty vector (EV) in the absence and presence (+dithio) of sodium dithionite were recorded on a spectrophotometer equipped with an integrating cavity. **B,** Difference spectra between hCYGB and EV cells in the absence (hCYGB) and presence of sodium dithionite. The maximum absorption at 417 nm in the Soret region and maxima at 540 and 578 nm are characteristic of oxycytoglobin (CygbFe^2+^-O_2_). **C,** Difference spectra between hCYGB and EV cells in the presence of increasing concentrations of hydrogen peroxide. For all experiments, each condition was performed at least three times on three independent days.

Since we did not observe any changes at the heme center consistent with reaction of hydrogen peroxide with cytoglobin, we monitored changes in oxidation states at the two surface cysteine residues on cytoglobin. For other proteins, this has been achieved by examining protein mobility in non-reducing SDS PAGE^32^. Following preparation of lysates from HEK cells expressing cytoglobin, recovery of cytoglobin in non-reducing gels was relatively low, presumably due to cytoglobin degradation or aggregation. We found that treatment of the samples with the thiol alkylating agent N-ethylmaleimide (NEM) or using the cytoglobin C83A and C38A variants increased recovery (**Figure 7A**), suggesting that this depended on either blockade of the two reactive cysteine residues by NEM or the absence of one cysteine residue. Significantly, following treatment of cytoglobin expressing cells with hydrogen peroxide or the thioredoxin reductase inhibitor auranofin, we also observed an increase in cytoglobin protein in non-reducing gels (**Figure 7B**). This was consistent with blockade of the cysteine residues through oxidation to intramolecular disulfides before preparation of the lysate. Overall, these results indicated intracellular oxidation of the cysteine surface residues by hydrogen peroxide or through inhibition of thioredoxin reductase.

**Figure 7.**
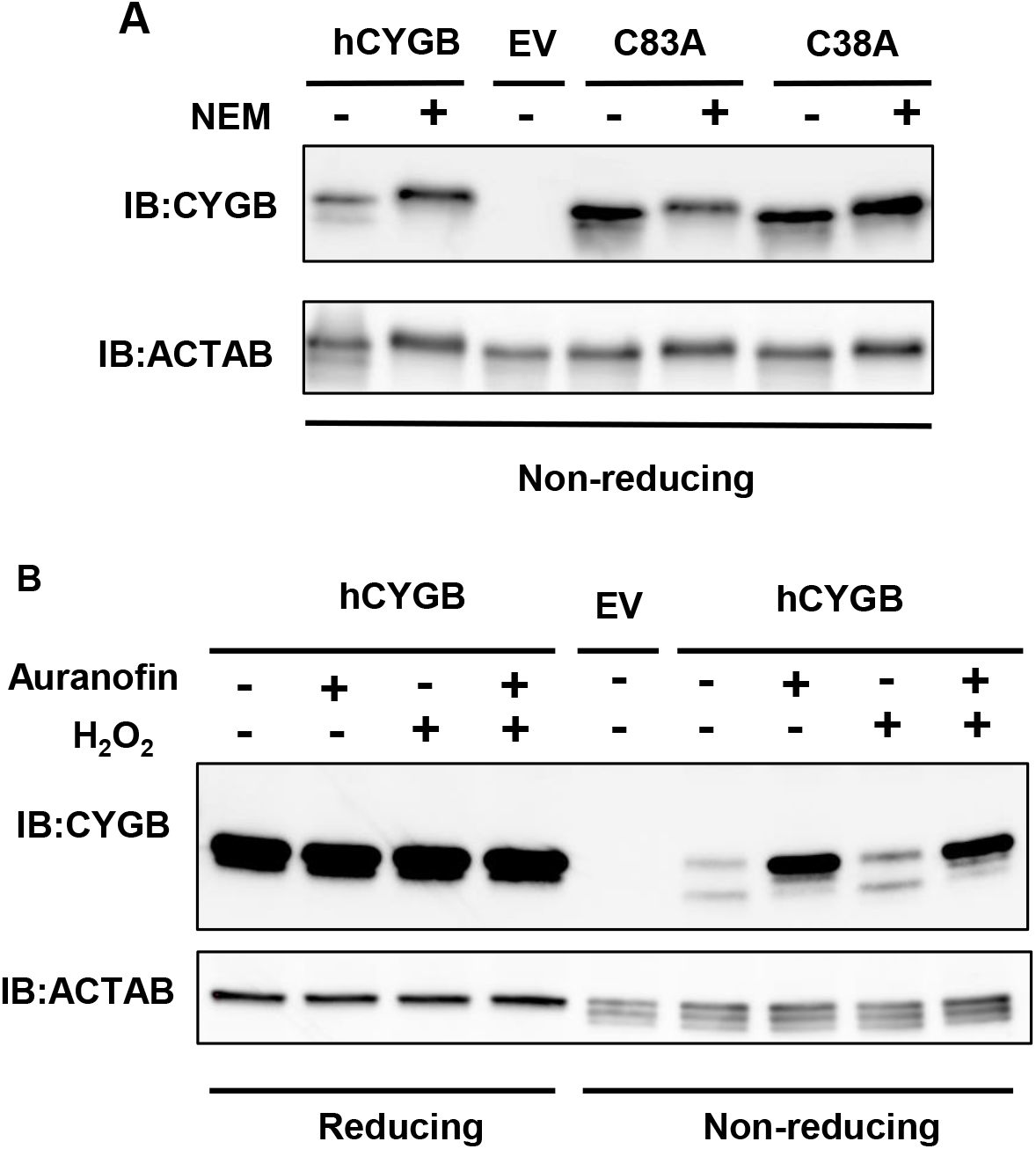
Intracellular oxidation of cytoglobin at cysteine residues. A, Cell lysates were prepared from HEK293 cells expressing an empty vector (EV), human cytoglobin (hCYGB), and C83A and C38A variants. Cells were incubated with or without NEM and cell lysates were prepared for SDS PAGE under non-reducing conditions following immunoblotting with an antibody against cytoglobin (IB:CYGB) with beta-actin (ACTAB) as an internal control. B, Immunoblots under reducing and non-reducing conditions of cell lysates prepared from HEK293 cells expressing an empty vector (EV) or human cytoglobin (hCYGB) and treated with 150 µM hydrogen peroxide for 10 minutes or auranofin for 30 min. All experiments have been repeated at least three times.

### 2.6 Loss of cytoglobin increases the sensitivity of glycolysis to hydrogen peroxide in isolated mouse carotid arteries

Cytoglobin protein expression has been described in rodent and medial vascular smooth muscle cells by our group and others^33,34^ and it has been implicated in the regulation of vascular tone and response to vascular injury ^34,35^. Even under fully aerobic conditions and in the absence of force generation, glycolysis provides a significant fraction of the basal bioenergetic requirement of vessels ^36^. Since we hypothesized that cytoglobin limits intracellular hydrogen peroxide, we predicted that deletion of cytoglobin in the mouse would result in the sensitization of glycolysis to hydrogen peroxide in vessels. We set out to test our hypothesis *ex vivo* in mouse carotid arteries by measuring extracellular acidification rates (ECAR) using Seahorse XF technology^37^.

Before proceeding with metabolic flux analyses on isolated mouse carotid arteries, we established the specific cellular association of cytoglobin in these vessels. Indirect immunostaining of cytoglobin protein in wild-type mice showed expression in the media consistent with SMC association, but also significant staining in the adventitia (**Figure 8A**). To confirm cellular association of cytoglobin with SMCs, we used transgenic mice that express a tamoxifen-inducible Cre-recombinase driven by the SMC-specific *Myh11* promoter combined with a Cre-responsive zsGreen inserted at the *ROSA26* locus. Using this system, SMCs are permanently labeled with zsGreen once the mice are administered tamoxifen. We stained carotid artery cross sections for cytoglobin by indirect immunofluorescence combined with *in situ* hybridization for dermatopontin transcripts (Dpt), a fibroblast marker^38^, cytoglobin co-associated with dermatopontin positive adventitial cells and ZsGreen positive medial cells (**Figure 8B**), indicating expression of cytoglobin in medial smooth muscle and adventitial fibroblasts in this particular vessel. Next, cytoglobin wild-type and knockout mice littermates (Cygb WT and Cygb KO mice) were generated (**Supplementary Figure 2**). Cytoglobin loss was verified by Western blot and indirect immunofluorescence analysis (**Supplementary Figure 2**). We performed a differential gene expression analysis on a bulk RNA seq from isolated left common carotid arteries (**Supplementary Figure 3**). We queried this data set for the expression of globins (**Figure 8C**) and established that cytoglobin mRNA exceeded the expression of any other globin transcripts by more than 10-fold. In addition, deletion of cytoglobin did not alter the expression of the other globin transcripts (**Figure 8C**). Finally, Gene Set Enrichment Analysis (GSEA) failed to yield any enrichment in genes associated with reactive oxygen species (**Supplementary Figure 3**).

**Fig. 8.**
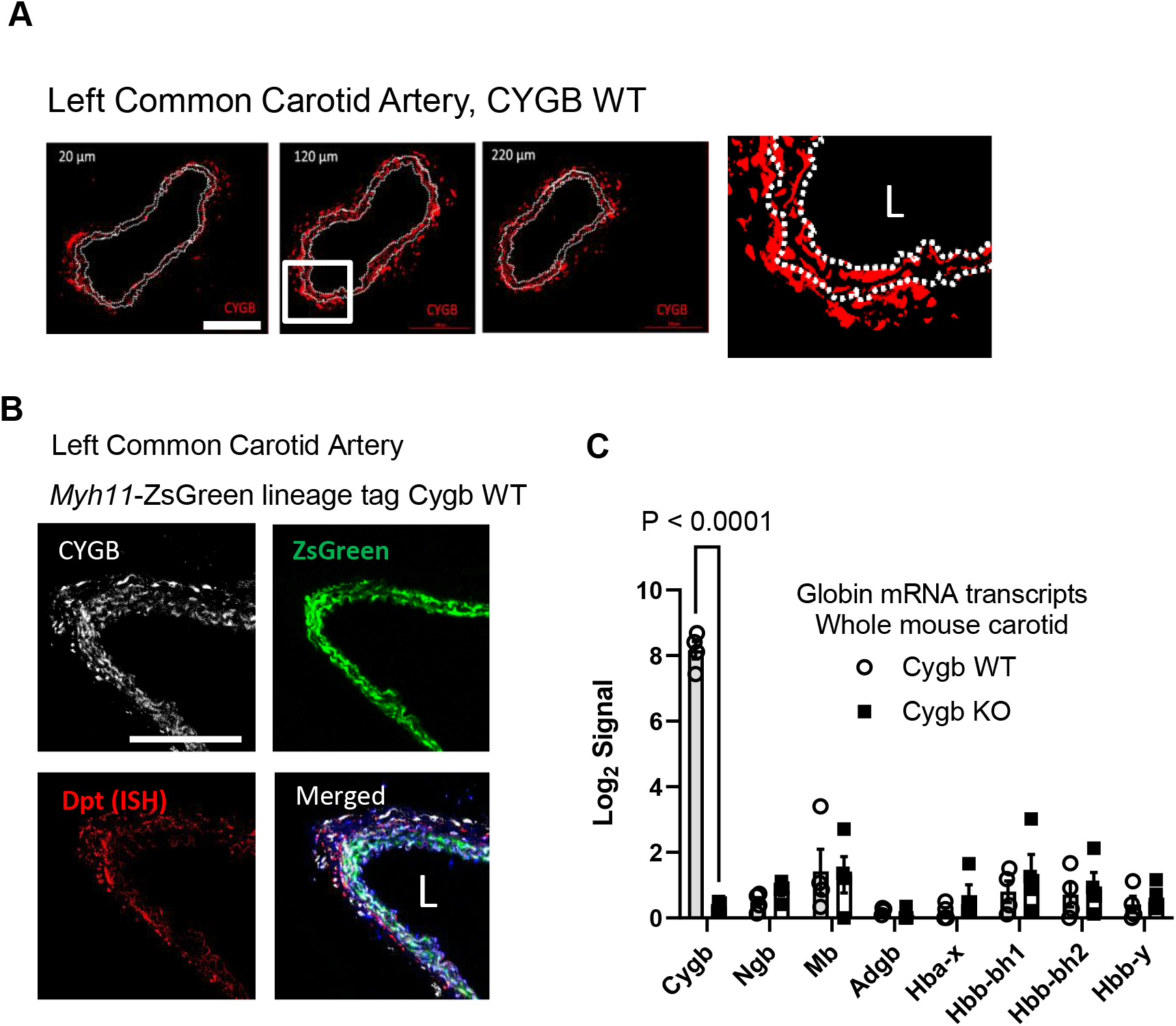
Cytoglobin expression in the mouse carotid artery. **A,** Representative indirect immunofluorescence staining for cytoglobin (red) in serial sections - 20, 120, and 220 µm from the bifurcation between the internal and external carotid artery - obtained from the left common artery of wildtype mice; scale bar is 200 µm; inset shows enlargement of the middle picture with cytoglobin staining in both the adventitia and media; white dotted line delineate the limit of the media, based on laminas autofluorescence. **B,** Representative immunofluorescence pictures showing indirect immunofluorescence for CYGB, ZsGreen positive cells, and in situ hybridization (ISH) for dermatopontin (Dpt) mRNA in mouse carotid artery from tamoxifen-induced Myh11 ER^T2^ cre ZsGreen mice. Cytoglobin is expressed in the medial smooth muscle cells and adventitial fibroblasts. Scale bar is 200 µm; L = lumen. **C,** Bulk RNA seq analysis was performed on the left common carotid artery of cytoglobin wildtype (Cygb WT) and knockout (Cygb KO) mouse littermates and results were queried for globin mRNA and plotted as the Log_2_ signal for each transcript. Cytoglobin is the primary globin expressed at the mRNA level in the mouse carotid artery of WT mice with no statistically significant changes in the other globins following deletion of cytoglobin. Statistical analysis was performed using two-way ANOVA.

Next, we isolated 2-mm length segments of mouse carotid arteries and measured basal ECAR following the sequential addition of a blank, the mitochondrial uncoupler FCCP and a combination of two respiratory chain inhibitors rotenone and antimycin A (**Figure 9A**). The proton ATPase inhibitor oligomycin was not used in these experiments due to lack of effect. Addition of FCCP in the presence of glucose increased ECAR suggesting that the basal glycolytic rate of these vessels is limited by a low resting demand that can be increased following an artificially high ATP demand through mitochondrial uncoupling with FCCP. The lack of effect of rotenone and antimycin A indicated the absence of contribution of oxidative phosphorylation to extracellular acidification. There was also no difference in basal ECAR or FCCP-stimulated ECAR between wild-type and knockout mice (**Supplementary Figure 4**). Treatment of the vessels with 150 μM hydrogen peroxide resulted in a decrease in ECAR (**Figure 9B and C**). However, the extent of the decrease in ECAR was statistically significantly greater in cytoglobin knockout vessels compared to wildtype (**Figure 9D**). We also noted a collapse in FCCP-stimulated ECAR (**Figure 9C and D**) and the vessels with cytoglobin deletion were unable to sustain a measurable increase, past a few minutes. Altogether, these results indicated a protective effect of cytoglobin against hydrogen peroxide-mediated inhibition of glycolysis in mouse carotid arteries.

**Fig. 9.**
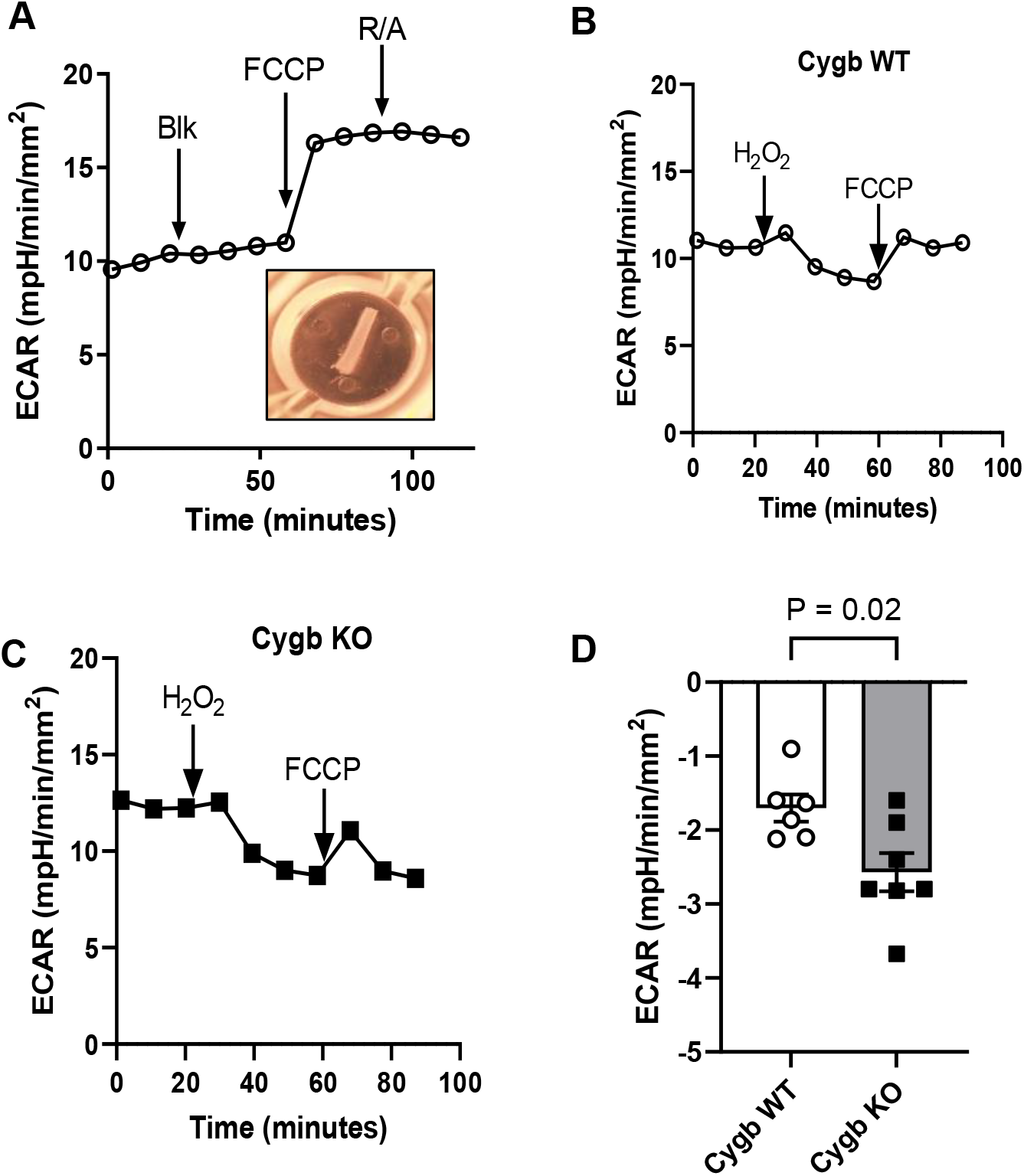
Loss of cytoglobin increases the sensitivity of glycolysis to hydrogen peroxide in isolated mouse carotid arteries. A, Representative tracing of extracellular acidification rate (ECAR) measurements obtained from the carotid artery of mice. Arrows indicate sequential addition of PBS (Blk), FCCP, and rotenone with antimycin A (R/A). Insert, representative image of an isolated length of the mouse carotid artery prepared for metabolic flux analysis. B, Extracellular acidification profiles obtained from carotid arteries from cytoglobin wildtype (Cygb WT, B) and knockout (Cygb KO, C) mouse littermates. Arrows indicate the sequential addition of hydrogen peroxide (H_2_O_2_, 200 μM) and FCCP. D, Quantitation of the decrease in ECAR following the addition of hydrogen peroxide on carotid arteries obtained from cytoglobin wildtype (Cygb WT) and knockout (Cygb KO) mice. Each data point indicates one mouse. Statistical analysis was performed using unpaired Student’s t-test.

## 3. Discussion

Cytoglobin can assume both hexa and pentacoordinated conformations to take on electron transfer, ligand binding, and peroxidase activities^9,30^. Biochemical studies also indicate relatively high rates of nitrite reduction, nitric oxide dioxygenation, and superoxide dismutation that all involve reactions at the heme center^5,39,40^. However, a clear understanding of cytoglobin physiological function is still lacking. In the present study, we used three independent cellular endpoints that are sensitive to hydrogen peroxide to specifically determine the role of cytoglobin as an intracellular scavenger of hydrogen peroxide.

The first question we asked was whether intracellular cytoglobin could compete for hydrogen peroxide with peroxiredoxin 1 and 2, the two primary thiol-based peroxidases that function in hydrogen peroxide detoxification in the cytosol^41^. These peroxiredoxins share similar structure and catalytic steps. However, peroxiredoxin 2 activity responds to lower levels of hydrogen peroxide than peroxiredoxin 1^42^. This agrees with our observation that a large fraction of peroxiredoxin 2 already existed as a dimer in control cells, while a sharp increase in peroxiredoxin 1 dimerization occurred only following the addition of exogenous hydrogen peroxide. The rapid loss of peroxiredoxin 2 dimers in the presence of exogenous hydrogen peroxide compared to peroxiredoxin 1 was also in agreement with the reported ten-fold slower rate of disulfide bond formation for peroxiredoxin 2^42,43^. This makes peroxiredoxin 2 more susceptible to hyperoxidation^33^, leading to the accumulation of its monomeric hyperoxidized inactive form. Thus, the persistence of peroxiredoxin 2 dimers and decrease in hyperoxidation following cytoglobin expression suggest a decrease in hydrogen peroxide load. However, the observation that cytoglobin did not affect peroxiredoxin 1 activity is more difficult to interpret. The protective effect of cytoglobin is likely to be more complex than simple competition for hydrogen peroxide between peroxiredoxin 1 and 2 and specific protein-protein interactions between cytoglobin and peroxiredoxin 2 might be required. For example, it is conceivable that the slower rate of disulfide bond formation for peroxiredoxin 2 may allow for cytoglobin to condense with peroxiredoxin 2 to form a mixed disulfide that may be reduced by the thioredoxin/thioredoxin reductase system. This issue will need to be addressed in future studies.

Remarkably, using the hydrogen peroxide sensor Hyper7-NES, we found that cytoglobin expression promoted a more reduced environment in HEK293 cells. The kinetics of Hyper7-NES oxidation and reduction following the addition of hydrogen peroxide also showed an increase in reduction rates of the probe by cytoglobin. It is important to understand that Hyper7 sensitivity to hydrogen peroxide relies on conformational changes induced by the reversible oxidation of two cysteine residues located on the probe to form a disulfide bridge^27^. Thus, the redox state of the cysteine pair and Hyper7 fluorescence depends not only on intracellular hydrogen peroxide concentrations but also on rates of disulfide reduction. Past studies have shown that Hyper7 is reduced by the thioredoxin/thioredoxin reductase system^27,29^. We found that the increase in reduction rates of Hyper7 by cytoglobin was lost after molecular or pharmacological inactivation of the thioredoxin and thioredoxin reductase. This indicated that cytoglobin antioxidant function relied on the same reductive pathway and did not recruit and was not reliant on alternate pathways.

The ability of cytoglobin to limit the inactivation of peroxiredoxin 2 and alter Hyper7 signal was still surprising to us because the heme peroxidase activity of cytoglobin is several folds slower than peroxiredoxin 2^9^. Thus, the next question was whether we could establish changes in cytoglobin heme oxidation following hydrogen peroxide challenge. We were able to directly determine the heme status of intracellular cytoglobin and demonstrated it existed primarily in the oxyferrous form. Although oxymyoglobin autoxidation is higher than other mammalian globins^44^, the strong reducing intracellular environment and presence of heme reductases would be predicted to limit oxidation of the heme center by hydrogen peroxide. In agreement, we found no evidence that high micromolar concentrations of exogenous hydrogen peroxide can oxidize the oxygenated heme of intracellular cytoglobin. A coordinated heme for its antioxidant activity is clearly required based on inhibition of reductive activity on Hyper7 following substitution of the proximal and distal histidines, but this might be necessary as a structural feature independent of heme oxidation.

Considering the lack of evidence of any oxidation at the heme center, we considered an alternative possibility. Cytoglobin contains two cysteine residues (C83 and C38) that may be oxidized to form an intramolecular disulfide bond. This has been shown to increase oxygen binding at the heme center by facilitating heme transition from its hexacoordinated to pentacoordinated conformation^45,46^. However, the physiological impact of these observations is unclear because even in the presence of reduced cysteine residues, the P_50_ of cytoglobin for oxygen is still very low (approximately 1 torr^45^). This dictates that cytoglobin is fully oxygenated at ambient oxygen tension in cell culture models, independent of the redox state of its cysteines. In contrast, the sensitivity of these cysteine residues to hydrogen peroxide has been recently highlighted *in vitro*^47^. Thus, we entertained the possibility that the antioxidant effect of cytoglobin may be explained through reaction of hydrogen peroxide with these cysteine residues instead of the heme center. Our studies that examined cytoglobin in non-reducing gels demonstrated the sensitivity of cytoglobin cysteine residues to hydrogen peroxide and pharmacological inhibition of thioredoxin reductase. However, we could not detect a change in cytoglobin protein mobility based on the reduction and oxidation states of the cysteine residues as shown for other proteins^32,48^. It might be possible to follow cytoglobin thiol oxidation to the disulfide using approaches like the kinetic trapping method developed by Schwetassek et al.^49^. However, we found that substitution of C83 only - and not C38 – reversed the effect of cytoglobin on Hyper7 reduction and GAPDH activity. This functional dichotomy between C83 and C38 is important. It would indicate alternative reactivities of the thiols for hydrogen peroxide and formation of oxidation states different than a disulfide bridge. Based on these results, we would like to propose that cytoglobin regulates hydrogen peroxide concentrations in cells primarily through alteration in the oxidation and reduction state of its cysteine residues. Detailed testing of this hypothesis will require *in vitro* experiments including reconstitution of cytoglobin with the redox systems that may catalyze cytoglobin antioxidant activity.

An essential short-term cellular signal in response to hydrogen peroxide is the rerouting of glycolytic fluxes to the oxidative branch of the pentose monophosphate pathway^31^. This increases the production of NADPH that is rapidly turned over by the higher demand for reducing equivalents from antioxidant enzymes. In this study, we show that cytoglobin regulates this response in cultured cells and in isolated vessels. Our finding that cytoglobin minimizes the sharp increase in G3P and F1,6P following hydrogen peroxide treatment that is mirrored by a decrease in ribose-5-phosphate again indicates a decrease in oxidant burden following cytoglobin expression. One mechanism that explains glycolytic rerouting is the reversible inhibition of GADPH by hydrogen peroxide at specific cysteine residues. The rate of reaction of GADPH with hydrogen peroxide is relatively slow (10^2^-10^3^ M^−1^.s^−1^)^50^. Assuming that the rate of reaction of cytoglobin cysteine residues is of the same order as GAPDH, cytoglobin might be able to regulate GAPDH through direct competitive reactions with hydrogen peroxide.

The role of cytoglobin in the cardiovascular system remains incompletely understood. Cytoglobin is expressed in medial vascular smooth muscle cells^33^ and we now show its expression in adventitial fibroblasts in the mouse carotid. Cytoglobin has been proposed to regulate nitric oxide-dependent vasoreactivity^51^ and studies from this laboratory have also shown that in two different rodent models of vascular injury, decrease in cytoglobin levels is associated with inhibition of neointima formation^34^. In addition, we found that the loss of cytoglobin sensitized human and rodent vascular smooth muscle cells to apoptosis and this was inhibited by the antioxidant N-acetyl cysteine^34^. While we had shown that loss of cytoglobin did not sensitize VSM cells to hydrogen peroxide cytotoxicity^34^, others had indicated increase in cytoglobin expression following hydrogen peroxide challenge, and protection against hydrogen peroxide following over-expression of cytoglobin^12,14,52,53^. We found that glycolytic fluxes in isolated mouse carotid arteries from global cytoglobin knockout mice were also more sensitive to hydrogen peroxide mediated inhibition. This is important because it indicates that the amount of tissue associated cytoglobin is sufficient to promote hydrogen peroxide decomposition and compete with other intracellular peroxidases. It is also possible that apparent reaction rates are increased from molecular co-association or co-localization of protein complexes and increase in local concentrations in subcellular compartments; however, this will need to be established.

In conclusion, we show that cytoglobin is an efficient scavenger of hydrogen peroxide in cells. We propose that cytoglobin decomposes hydrogen peroxide through reaction at cysteine residues rather than at the heme center. However, it will be important to clarify the role of the heme center as a possible regulatory site through ligand binding or reaction to produce oxidants such as hydrogen peroxide^5^. We expect that further understanding of the mechanism of action of intracellular cytoglobin, combined with *in vitro* reconstitution experiments to elucidate the reduction system involved, will allow to manipulate the physiological function of cytoglobin and integrate this new knowledge to understand human diseases.

## 4. Materials and methods

### 4.1 Supplies and reagents

All supplies and reagents are listed in Supplementary Table 1.

### 4.2 Mice

All procedures related to mice were approved by the Institutional and Animal Care Use Committee at Albany Medical College. The cytoglobin “knock out first” allele was obtained from the University of Toronto, Canada, and global wild-type and cytoglobin knockout mice littermates (Cygb WT mice and Cygb KO) were generated from heterozygous breeding pairs. Both males and females were used and randomly assigned to experimental groups. Myh11-Cre^ERT^ was a kind gift from Dr. Singer (Albany Medical Center) and ROSA26-zsGreen reporter mice were obtained from the Jackson Laboratory. Myh11-Cre^ERT^ transgenic mice and ROSA26-zsGreen were bred to generate tamoxifen-inducible SMC-specific zsGreen reporter mice. Mice were housed in pathogen-free rooms with controlled light-dark cycle, temperature, and humidity. Mice were kept in cages in groups of five or fewer with ad libitum access to food and water. Eight-to thirteen-week-old mice were euthanized and the thoracic aorta, left common carotid artery, heart, liver and spleen were collected for immunoblotting, imaging, and metabolic flux studies.

### 4.3 Transcriptomic analysis

RNA quantity was determined using a Qubit Flex Fluorometer. Library preparation was performed using an Ion Chef System followed by sequencing using an Ion GeneStudio S5 Plus System (All from Thermo Fisher Scientific, San Jose, CA). We used the Ion AmpliSeq Transcriptome Mouse Gene Expression Kit for mouse the left common carotid artery and the Ion AmpliSeq Transcriptome Human Gene Expression Kit for HEK293 cells.

### 4.4 Generation of HEK 293 Clones, cell culture, and treatment with hydrogen peroxide

A pcDNA 3.1 plasmid containing the full length human cytoglobin gene was purchased from GenScript. The empty vector control and specific point mutations of this plasmid were also generated by GenScript. Plasmids were transformed with Invitrogen’s One Shot Top10 chemically competent cells, streaked onto LB agar plates with ampicillin selection and incubated overnight at 37°C to achieve individual colonies. LB broth containing 100 µg/ml Ampicillin was inoculated with one of the resulting colonies and incubated at 37°C with shaking overnight. Plasmid DNA was recovered from the LB broth with a NucleoBond Xtra Midi plus kit as per their protocol. HEK293 cells were transfected with Dharmafect kb, stably expressing human Cytoglobin cell lines were established through limiting dilution cloning. Cells were maintained in DMEM supplemented with 10% fetal calf serum, 2 mM L-glutamine, and Geneticin for selection.

For most experiments using adherent HEK 293 clones, one million cells were seeded in 60 mm dishes with 10% DMEM + 750 µg/ml Geneticin to a final volume of 4 mls. Forty-eight hours post-seeding the cells were treated with 20 to 200 µM H_2_O_2_ for 2 to 20 minutes.

### 4.5 Spectrophotometric study of intracellular cytoglobin

Cytoglobin quantitation in HEK293 clones was performed as previously described with some modifications^38^. Briefly, absorbances of whole cell suspensions were measured using an Olis-Clarity VF spectrophotometer using an 8-ml cuvette at a final concentration varying from 1×10^6^ to 1×10^7^ cells/ml. Apparent absorbance values were recorded relative to a buffer baseline. Live cell samples were read before and after the addition of dithionite. Spectra subtraction between cells with empty vector and cells expressing cytoglobin was performed using Olis software and plotted using Graph Pad Prism v9.

### 4.6 Reducing and non-reducing immunoblotting

Following treatment with hydrogen peroxide for the specified time, 50 mM NEM was added to each plate and incubated for 2 minutes at RT. Media was aspirated and cells were washed with PBS containing 50 mM NEM. Cells were lysed in 500 µl of RIPA buffer containing 50 mM NEM and HALT protease/phosphatase inhibitors (1:100). Lysates were spun at 12000 × g for 20 min at 4 °C. Supernatants were transferred into fresh tubes and protein concentration was determined using Pierce BCA protein assay. Samples were divided into two; one received an equal volume of 2X SDS sample buffer with 720 mM β-mercaptoethanol (reducing) and the other 2X SDS-sample buffer without β-mercaptoethanol (non-reducing); samples were heated to 95°C for 5 minutes. Equal amounts of total protein were loaded on a 4-20% gradient gel and electrophoresed until the dye front reached the bottom of the gel. Proteins were transferred to PVDF membranes at 100V for 1 hour. Membranes were blocked overnight with 5% NFM/TBST at 4°C. Membranes were incubated with primary antibodies for peroxiredoxin 1 (1:10000) or peroxiredoxin 2 (1:10000) for 1 hour at room temperature, washed 3 × 15 minutes, incubated with anti-mouse-HRP/anti Rabbit-HRP secondary (1:2500) for 1 hour at RT, washed 3 × 15 minutes, then visualized using BioRad Clarity Western ECL substrate on a BioRad ChemiDoc MP imaging system.

### 4.7 Multiplex immunoblotting

Cell lysates were prepared in RIPA/HALT buffer as outlined above. Protein was mixed with reducing SDS sample buffer, heated to 95°C for 5 minutes and loaded into wells of 4-20% gradient gels at 15 ug/lane. The gels were run at 100V until the dye fronts reached the bottom; proteins were transferred to low Fluorescence PVDF membranes and blocked in 5% non-fat milk/TBST overnight. Membranes were incubated with anti-rabbit Cytoglobin pAb (1:2000) for 1 hour at room temperature, washed 3 × 15 min in TBST. Blots were incubated in a secondary fluorescent goat anti rabbit StarBright blue 700 (1:3000), and anti-Actin hFAB Rho (1:3000) for 1 hour at room temperature, blots were washed 3 × 15 min in TBST, allowed to dry and imaged with the BioRad ChemiDoc MP imaging system using the Multi Red Blue and Green program.

### 4.8 Metabolic flux studies and GAPDH activity

*Ex vivo* studies of metabolic fluxes in isolated vessels were performed as previously described with some modifications^37^. Briefly, the left common carotid arteries were collected from 8-13 weeks old CYGB WT and KO mouse littermates after perfusion with PBS under isoflurane anesthesia. The carotid arteries were cleaned of exterior fat under a dissecting microscope and cut into two-millimeter sections. Individual wells of XF96 culture plates precoated with CELL-TAK were loaded with 180 µl of XF Base Media containing 10 mM glucose pH 7.4. Plates were equilibrated for 1 hour at 37 L in an incubator without CO_2_ prior to the addition of the 2 mm carotid artery sections, the plates were briefly spun at 500 rpm for 1 minute to adhere vessels to the bottom of the wells. Test compounds were loaded into the corresponding ports of the Sensor cartridge prior to instrument calibration. Acidification rates were measured on a SeaHorse XFe96 analyzer. Titration experiments were performed to establish the optimal dosage hydrogen peroxide (250 µM), FCCP (1 µM), and the rotenone/antimycin A mixture (0.5 µM). We found that oligomycin did not provide reproducible results, most likely due to limited penetrance. Each extracellular acidification rate measurement included 3 min of mixing, 3 min wait time, and 3 min of continuous measurement with at least three measurement rounds per injection. Following the measurements, images of each vessel were taken for determination of vessel length to normalize ECAR data to vessel surface area. For each experiment, at least two 2-mm pieces were used per mouse and averaged. Biological replicates were individual mice and both males and females were used for these experiments.

### 4.9 Metabolomics mass spectrometry analysis

Methods were performed as described previously^54^. Briefly, snap frozen cell pellet samples were kept in the −80 °C freezer until time of extraction. To extract metabolites, samples were extracted with 1 mL of a chilled 40:40:20 Acetonitrile:Methanol:Water. Samples were lysed by sonication for 20 seconds at 4 °C, then centrifuged at 4 °C for 2 minutes at a speed of 14000 × g. Then 200 µL of the supernatant from each tube was transferred into a low volume amber borosilicate glass autosampler vial with tapered insert and dried by vacuum concentrator. Samples were resuspended in 100 µL of water for LC-MS/MS analysis. A mixture of standards was prepared at 12 concentrations and used for quantification.

For each sample, 10 µL was loaded onto an IonPac AS-11 HC strong anion-exchange analytical column (2 mm × 250 mm, 4 µm particle diameter, Thermo Scientific) with AG-11 HC guard column (2 mm × 50 mm) heated to 40 °C. An Ultimate (Thermo Scientific) LC system was used for applying a 0.35 mL/min gradient of mobile phase A (Water, degassed) and mobile phase B (100 mM sodium hydroxide, degassed): hold 12.5% B for 5 min, increase to 20% over 5 min, increase to 27.5 B over 7.5 min, increase to 42.5% B over 5 min increase to 95% B over 10 min, hold at 95% B for 9.5 min, then returning to 12.5% B over 0.5 min. Post column eluent was passed through a 2 mm AERS 500e suppressor operated via a reagent-free controller (RFC-10, Dionex) set to 50 mA, a binary pump (Agilent 1100) was used to deliver water at 0.5 mL/min for external water regeneration mode. Then eluent was ionized by electrospray ionization to be analyzed by a TSQ Quantiva mass spectrometer (Thermo Scientific). The following source settings were used: 350°C vaporization temperature, 350°C ion transfer tube temperature, 18 units Sheath Gas, 4 unit Aux Gas, 1 unit Sweep gas, polarity switching |3800 V| (1.5-45 min), and Use Calibrated RF Lens option. MS was operated in single reaction monitoring mode with specified transitions, retention time windows and colliisions energies, using 0.7 FWHM resolution for Q1 and 1.2 FWHM for Q3, 0.8 s cycle time, 1.5 mTorr CID gas, and 30 s Chrom Filter.

Raw files were processed using Xcalibur Quan Browser (v4.0.27.10, Thermo Scientific); The prepared mixture of standards was used to locate appropriate peaks for peak areas analysis.

### 4.11 Real-time imaging

HEK 293 cells expressing human cytoglobin or empty vector were seeded into MatTek 35 mm glass bottom culture dishes in antibiotic free DMEM supplemented with 10% Fetal Calf Serum and 2 mM L-Glutamine. Cells were transfected with the pCS2+Hyper7 plasmid using Dharmafect kb transfection reagent according to the manufacturer’s recommendations. Twenty-four hours post-transfection, culture medium was replaced with 1.2 mL of HBSS supplemented with 20 mM HEPES. For some experiments, cells were pre-treated with10 µM final concentration auranofin for 30 minutes prior to HyPer7 analysis. For other experiments, cells were transiently transfected with small interfering RNA (siRNA) oligonucleotide against thioredoxin 1 or nontarget region using Dharmafect 1. Cells were incubated in Opti-MEM and supplemented with 5% fetal bovine serum (FBS) for 72 hours following transfection. Cells were subsequently transfected with HyPer 7 NES as described above.

Cell imaging was performed using a Leica DMI 8 Thunder microscope, equipped with a HC PL APO 63x 1.4NA oil objective at 37 °C. Samples were excited sequentially via 440/15 and 510/15 band-pass excitation filters. Emission was collected every 20 seconds using a 519/25 bandpass emission filter. After 5–10 images were acquired, a small volume of hydrogen peroxide was carefully added to obtain a final concentration of 200 µM. The time series were analyzed via Fiji freeware (https://fiji.sc). The background was subtracted from 440 and 510 nm stacks. Every 510 nm stack was divided by the corresponding 440 nm stack frame-by-frame. The resulting stack was depicted in pseudo-colors using a “16-colors” lookup table. The plasmid pCS2+Hyper7-NES was a gift from Dr. Belousov (Addgene plasmid # 136467).

### 4.12 Immunofluorescence

For tissues, 10 µm tissue sections were cut from prepared OCT blocks using a Leica CM1850 cryostat and transferred to charged microscope slides, allowed to dry at room temperature and then stored at −80°c until use. For immunofluorescence, the sections were removed from the freezer, air dried, fixed with ice cold acetone for 10 minutes air dried again, then outlined with a hydrophobic barrier pen. Tissues were rehydrated with PBS, blocked with 5% sera representing the secondary antibody species for at least 1 hour; rinsed with PBS/0.1% TX 100 (PBST). Tissues were incubated with anti-rabbit cytoglobin polyclonal antibody (1:100) with a matching rabbit IgG isotype control for 1 hour at room temperature, washed again in PBST, incubated with goat anti rabbit Alexafluor 594 secondary antibody (1:200) for 1 hour at room temperature. Slides were then washed once with PBST, once with a 50:50 PBS/water mix; incubated with DAPI for 15 minutes at room temperature, then rinsed again in the 50:50 mixture; cover slipped with VectaShield Antifade Mounting Medium; sealed with clear nail polish and stored protected from light at 4°C.

For cellular immunofluorescence studies, cells were plated in 8 well glass bottom Ibidi slides precoated with poly-L-lysine in DMEM supplemented with 10% FBS and 2 mM L-glutamine then returned to 37°C CO_2_ incubator to attach overnight. Cells were fixed in 4% formaldehyde for 15 minutes at room temperature washed with PBS then permeabilized in PBS/0.2% Triton-X100 for 5 minutes at room temperature, blocked with 5% goat serum for at least 1 hr. After one rinse with PBS/0.1% Triton-X100 (PBST), cells were incubated with a polyclonal rabbit anti-cytoglobin (1:100) for 1 hour at room temperature, washed again in PBST, incubated with Goat anti Rabbit AlexaFluor 594 secondary antibody (1:200) for 1 hour at room temperature, washed once with PBST, once with a 50:50 Water/PBS mix then stained with DAPI for 15 minutes. After one final wash in the 50:50 mixture the slides were cover slipped using VectaShield Antifade Mounting media, sealed with clear nail polish and stored at 4oC in the dark until imaging.

Fluorescence in situ hybridization staining for dermatopontin transcripts was performed using Advanced Cell Diagnostics RNAscope™ Multiplex Fluorescent V2 Assay and the standard protocol with the Mm-Dpt probe along with the positive control probe Mm-Ppib and the negative control probe DapB. After the *in-situ* hybridization protocol, sections were stained for cytoglobin as described above.

### 4.13 Statistical analysis

Statistical analyses were performed with GraphPad Prism 9.0. The specific statistical test used to analyze each data set is specified in individual figure legends and p-values are shown in figures. A p-value less than 0.05 was considered statistically significant.

## Acknowledgments

This work was supported by NIH grants RO1 HL142807 (to D.J.), P41 GM108538 (to J.C.C.), RO1 HL13070 (to A.J.), and RO1 HL160661 (to A.J.).

## References

1. Schechter, A.N. Hemoglobin research and the origins of molecular medicine. Blood 112, 3927–3938 (2008).

2. Kawada, N., et al. Characterization of a stellate cell activation-associated protein (STAP) with peroxidase activity found in rat hepatic stellate cells. J Biol Chem 276, 25318–25323 (2001).

3. Trent, J.T., 3rd & Hargrove, M.S. A ubiquitously expressed human hexacoordinate hemoglobin. J Biol Chem 277, 19538–19545 (2002).

4. Burmester, T., Ebner, B., Weich, B. & Hankeln, T. Cytoglobin: a novel globin type ubiquitously expressed in vertebrate tissues. Molecular biology and evolution 19, 416–421 (2002).

5. Zweier, J.L., et al. Cytoglobin has potent superoxide dismutase function. Proc Natl Acad Sci U S A 118(2021).

6. Beckerson, P., Svistunenko, D. & Reeder, B. Effect of the distal histidine on the peroxidatic activity of monomeric cytoglobin. F1000Res 4, 87 (2015).

7. Reeder, B.J., Svistunenko, D.A. & Wilson, M.T. Lipid binding to cytoglobin leads to a change in haem co-ordination: a role for cytoglobin in lipid signalling of oxidative stress. The Biochemical journal 434, 483–492 (2011).

8. Tejero, J., et al. Peroxidase activation of cytoglobin by anionic phospholipids: Mechanisms and consequences. Biochim Biophys Acta 1861, 391–401 (2016).

9. Mathai, C., Jourd’heuil, F.L., Lopez-Soler, R.I. & Jourd’heuil, D. Emerging perspectives on cytoglobin, beyond NO dioxygenase and peroxidase. Redox Biol 32, 101468 (2020).

10. Vlasova, II. Peroxidase Activity of Human Hemoproteins: Keeping the Fire under Control. Molecules 23(2018).

11. Fordel, E., Geuens, E., Dewilde, S., De Coen, W. & Moens, L. Hypoxia/ischemia and the regulation of neuroglobin and cytoglobin expression. IUBMB Life 56, 681–687 (2004).

12. Li, D., et al. Cytoglobin up-regulated by hydrogen peroxide plays a protective role in oxidative stress. Neurochemical research 32, 1375–1380 (2007).

13. Xu, R., et al. Cytoglobin overexpression protects against damage-induced fibrosis. Mol Ther 13, 1093–1100 (2006).

14. Fordel, E., et al. Neuroglobin and cytoglobin overexpression protects human SH-SY5Y neuroblastoma cells against oxidative stress-induced cell death. Neurosci Lett 410, 146–151 (2006).

15. McRonald, F.E., Risk, J.M. & Hodges, N.J. Protection from intracellular oxidative stress by cytoglobin in normal and cancerous oesophageal cells. PloS one 7, e30587 (2012).

16. Thuy le, T.T., et al. Promotion of liver and lung tumorigenesis in DEN-treated cytoglobin-deficient mice. The American journal of pathology 179, 1050–1060 (2011).

17. Singh, S., et al. Cytoglobin modulates myogenic progenitor cell viability and muscle regeneration. Proc Natl Acad Sci U S A 111, E129–138 (2014).

18. Thuy le, T.T., et al. Absence of cytoglobin promotes multiple organ abnormalities in aged mice. Sci Rep 6, 24990 (2016).

19. Zhang, S., et al. Cytoglobin Promotes Cardiac Progenitor Cell Survival against Oxidative Stress via the Upregulation of the NFkappaB/iNOS Signal Pathway and Nitric Oxide Production. Sci Rep 7, 10754 (2017).

20. Cox, A.G., Winterbourn, C.C. & Hampton, M.B. Measuring the redox state of cellular peroxiredoxins by immunoblotting. Methods Enzymol 474, 51–66 (2010).

21. Karplus, P.A. A primer on peroxiredoxin biochemistry. Free radical biology & medicine 80, 183–190 (2015).

22. Peskin, A.V., et al. Hyperoxidation of peroxiredoxins 2 and 3: rate constants for the reactions of the sulfenic acid of the peroxidatic cysteine. J Biol Chem 288, 14170–14177 (2013).

23. Haynes, A.C., Qian, J., Reisz, J.A., Furdui, C.M. & Lowther, W.T. Molecular basis for the resistance of human mitochondrial 2-Cys peroxiredoxin 3 to hyperoxidation. J Biol Chem 288, 29714–29723 (2013).

24. Yang, K.S., et al. Inactivation of human peroxiredoxin I during catalysis as the result of the oxidation of the catalytic site cysteine to cysteine-sulfinic acid. J Biol Chem 277, 38029–38036 (2002).

25. Woo, H.A., et al. Reversible oxidation of the active site cysteine of peroxiredoxins to cysteine sulfinic acid. Immunoblot detection with antibodies specific for the hyperoxidized cysteine-containing sequence. J Biol Chem 278, 47361–47364 (2003).

26. Low, F.M., Hampton, M.B., Peskin, A.V. & Winterbourn, C.C. Peroxiredoxin 2 functions as a noncatalytic scavenger of low-level hydrogen peroxide in the erythrocyte. Blood 109, 2611–2617 (2007).

27. Pak, V.V., et al. Ultrasensitive Genetically Encoded Indicator for Hydrogen Peroxide Identifies Roles for the Oxidant in Cell Migration and Mitochondrial Function. Cell Metab 31, 642–653 e646 (2020).

28. Hoehne, M.N., et al. Spatial and temporal control of mitochondrial H(2) O(2) release in intact human cells. EMBO J 41, e109169 (2022).

29. de Cubas, L., Pak, V.V., Belousov, V.V., Ayte, J. & Hidalgo, E. The Mitochondria-to-Cytosol H(2)O(2) Gradient Is Caused by Peroxiredoxin-Dependent Cytosolic Scavenging. Antioxidants (Basel) 10(2021).

30. Kakar, S., Hoffman, F.G., Storz, J.F., Fabian, M. & Hargrove, M.S. Structure and reactivity of hexacoordinate hemoglobins. Biophys Chem 152, 1–14 (2010).

31. Kuehne, A., et al. Acute Activation of Oxidative Pentose Phosphate Pathway as First-Line Response to Oxidative Stress in Human Skin Cells. Molecular cell 59, 359–371 (2015).

32. Kwak, M.S., et al. Peroxiredoxin-mediated disulfide bond formation is required for nucleocytoplasmic translocation and secretion of HMGB1 in response to inflammatory stimuli. Redox Biol 24, 101203 (2019).

33. Halligan, K.E., Jourd’heuil, F.L. & Jourd’heuil, D. Cytoglobin is expressed in the vasculature and regulates cell respiration and proliferation via nitric oxide dioxygenation. J Biol Chem 284, 8539–8547 (2009).

34. Jourd’heuil, F.L., et al. The Hemoglobin Homolog Cytoglobin in Smooth Muscle Inhibits Apoptosis and Regulates Vascular Remodeling. Arterioscler Thromb Vasc Biol (2017).

35. Zweier, J. & Ilangovan, G. Regulation of Nitric Oxide Metabolism and Vascular Tone by Cytoglobin. Antioxid Redox Signal (2019).

36. Paul, R.J. Smooth muscle energetics. Annu Rev Physiol 51, 331–349 (1989).

37. Feeley, K.P., Westbrook, D.G., Bray, A.W. & Ballinger, S.W. An ex-vivo model for evaluating bioenergetics in aortic rings. Redox Biol 2, 1003–1007 (2014).

38. Buechler, M.B., et al. Cross-tissue organization of the fibroblast lineage. Nature 593, 575–579 (2021).

39. Petersen, M.G., Dewilde, S. & Fago, A. Reactions of ferrous neuroglobin and cytoglobin with nitrite under anaerobic conditions. J Inorg Biochem 102, 1777–1782 (2008).

40. Smagghe, B.J., Trent, J.T., 3rd & Hargrove, M.S. NO dioxygenase activity in hemoglobins is ubiquitous in vitro, but limited by reduction in vivo. PloS one 3, e2039 (2008).

41. Perkins, A., Poole, L.B. & Karplus, P.A. Tuning of peroxiredoxin catalysis for various physiological roles. Biochemistry 53, 7693–7705 (2014).

42. Portillo-Ledesma, S., et al. Differential Kinetics of Two-Cysteine Peroxiredoxin Disulfide Formation Reveal a Novel Model for Peroxide Sensing. Biochemistry 57, 3416–3424 (2018).

43. Dalla Rizza, J., Randall, L.M., Santos, J., Ferrer-Sueta, G. & Denicola, A. Differential parameters between cytosolic 2-Cys peroxiredoxins, PRDX1 and PRDX2. Protein Sci 28, 191–201 (2019).

44. Kiger, L., et al. Electron transfer function versus oxygen delivery: a comparative study for several hexacoordinated globins across the animal kingdom. PloS one 6, e20478 (2011).

45. Pesce, A., et al. Reversible hexa-to penta-coordination of the heme Fe atom modulates ligand binding properties of neuroglobin and cytoglobin. IUBMB Life 56, 657–664 (2004).

46. Hamdane, D., et al. The redox state of the cell regulates the ligand binding affinity of human neuroglobin and cytoglobin. J Biol Chem 278, 51713–51721 (2003).

47. DeMartino, A.W., et al. Redox sensor properties of human cytoglobin allosterically regulate heme pocket reactivity. Free radical biology & medicine 162, 423–434 (2021).

48. Simoes, V., et al. Redox-sensitive E2 Rad6 controls cellular response to oxidative stress via K63-linked ubiquitination of ribosomes. Cell Rep 39, 110860 (2022).

49. Schwertassek, U., et al. Selective redox regulation of cytokine receptor signaling by extracellular thioredoxin-1. EMBO J 26, 3086–3097 (2007).

50. Winterbourn, C.C. & Hampton, M.B. Thiol chemistry and specificity in redox signaling. Free radical biology & medicine 45, 549–561 (2008).

51. Liu, X., et al. Cytoglobin regulates blood pressure and vascular tone through nitric oxide metabolism in the vascular wall. Nat Commun 8, 14807 (2017).

52. Nishi, H., et al. Cytoglobin, a novel member of the globin family, protects kidney fibroblasts against oxidative stress under ischemic conditions. The American journal of pathology 178, 128–139 (2011).

53. Hoang, D.V., et al. Cytoglobin attenuates pancreatic cancer growth via scavenging reactive oxygen species. Oncogenesis 11, 23 (2022).

54. Overmyer, K.A., et al. Large-Scale Multi-omic Analysis of COVID-19 Severity. Cell Syst 12, 23–40 e27 (2021).

