## Supplemental Material for "A role for cytoglobin in regulating intracellular hydrogen peroxide and redox signals in the vasculature"

**SUPPLEMENTARY TABLE 1**

**SUPPLEMENTARY FIGURES**

**A role for cytoglobin in regulating intracellular hydrogen peroxide and redox signals in the vasculature**

Clinton Mathai^1^, Frances Jourd’heuil^1^, Le Gia Cat Pham^1^, Kurrim Gilliard^1^, Joseph Balnis^1, 2^, Annie Jen^3^, Katherine A. Overmyer^3,4^, Joshua J Coon^3,4,5^, Ariel Jaitovich^1,2^, Benoit Boivin^6^, and David Jourd’heuil^1^

^1^Department of Molecular and Cellular Physiology, Albany Medical College, Albany, NY

^2^Division of Pulmonary and Critical Care Medicine, Albany Medical College, Albany, NY

^3^Department of Biomolecular Chemistry, University of Wisconsin-Madison, Madison, WI

^4^Morgridge Institute for Research, Madison, WI, USA

^5^Department of Chemistry, University of Wisconsin-Madison, Madison, WI, USA

^6^College of Nanoscale Science & Engineering, SUNY Polytechnic Institute, Albany, NY, USA

**Supplementary Table 1: Reagents and Supplies**

| **Cells/Plasmids/siRNA** | **Supplier** | **Cat#** |
| --- | --- | --- |
| Cells HEK 293 | ATCC | CRL-1573 |
| DharmaFect 1 (transfection reagent) | Horizon Discovery | T-2001-02 |
| DharmaFect kb (transfection reagent) | Horizon Discovery | T-2006-01 |
| ON-TARGETplus Human TXN siRNA | Horizon Discovery | L-006340-00-005 |
| ON-Targetplus non targeting control pool | Horizon Discovery | D-001810-50 |
| pcDNA 3.1 (empty vector) | Genscript |  |
| pcDNA 3.1-hCYGB | Genscript | OHu14114C |
| pCS2+HyPer7 | Addgene | 136466 |
| pCS2+HyPer7-NES | Addgene | 136467 |
| pCS2+HyPer7-NLS | Addgene | 136468 |
| **Reagents** |  |  |
| 37% formaldehyde solution | Sigma | 252549-100 |
| Acetone | Sigma | 179124-1L |
| Acetonitrile Optima LC/MS | Thermo Scientific | A955-1 |
| Ampicillin sodium salt | Sigma | A0166 |
| Antimycin A | Sigma | A8674 |
| Auranofin | Enzo | BML-E1206-0100 |
| Bacto Tryptone | BD | 211705 |
| Bacto Yeast extract | BD | 212750 |
| BCA Protein Assay kit (Pierce) | Thermo Scientific | 23225 |
| bME (2-Mercaptoethanol) | Simga | 63689 |
| Cell-Tak tissue adhesive | Corning | 354240 |
| Clairty Western ECL Substrate | BioRad | 1705061 |
| D-(+)glucose solution | Sigma | G8769 |
| DAPI | Sigma | D9542-5mg |
| Difco Agar | BD | 214530 |
| DMEM (high Glucose) | Corning | 10-031-CV |
| DPBS | Corning | 21-031-CV |
| FBS | Cytiva | SH30396-03 |
| FCCP (Carbonyl cyanide 4-(trifluoromeothoxy)phenylhydarzone) | Sigma | C2920 |
| Geneticin (G418 sulfate) | Gibco | 10131-035 |
| HALT protease/phosphatase inhibitor | Thermo Scientific | 1861284 |
| HBSS | Corning | 21-031-CV |
| HBSS with calcium and magnesium | Corning | 21-023-CV |
| HEPES | Sigma | H4034-100 |
| Hydrogen Peroxide | Sigma | 216763 |
| Laemmli Sample buffer 2X | BioRad | 161-0737 |
| Laemmli Sample buffer 4X | BioRad | 161-0704 |
| L-Glutamine 200 mM | Sigma | G7513 |
| Methanol Optima LC/MS | Thermo Scientific | A456-1 |
| Molecular weigh markers (Percision Plus Western C) | BioRad | 161-0385 |
| NaCl | Sigma | S7653 |
| NEM (N-Ethylmaleimide) | Sigma | 04259 |
| Non-Fat Milk powder | Fisher | 50-751-7665 |
| OCT compound | Tissue-Tek | 4583 |
| Oligomycin | Sigma | O4876 |
| OneShot Top10 chemically competent cells | Invitrogen | C404003 |
| Optimem | Gibco | 31985-062 |
| Poly-L-lysine solution | Sigma | P4832 |
| RIPA buffer | Sigma | R0278 |
| Rotenone | Sigma | R8857 |
| Seahourse XF Base Medium | Agilent | 102353 |
| Sodium Hydrosulfite (dithionite) | Sigma | 157953 |
| Sodium Hydroxide | Thermo Scientific | O41281.06 |
| Sun Flower oil | Sigma | S5007 |
| Tamoxifen | Sigma | T5648 |
| Triton X-100 | Sigma | T9284 |
| Tween 20 | Sigma | P2287 |
| Vectashield Vibrance mounting media | Vector labs | H-1700 |
| Water Optima LC/MS | Thermo Scientific | W6-1 |
| XF-Base media | Agilent | 102353-100 |
| **Kits/Materials** |  |  |
| 8 Well glass chamber slides | Ibidi | 80841 |
| 35 mm round glass bottom | Mattek Corp | P35G-1.5-14C |
| 4-20% Mini Protean TGX stain Free gels | BioRad | 4568094 |
| Colorfrost plus microscope slides | Cardinal health | M6148-3P |
| GAPDH Activity kit | Sigma | MAL277 |
| Ion AmpliSeq Transcriptome Human Gene Expression Kit | Thermo Scientific | A26326 |
| Ion AmpliSeq Transcriptome Mouse Gene Expression Kit | Thermo Scientific | A3655A |
| Low Fluorescent PVDF Membrane | BioRad | 20190508 |
| Negative control probe DapB | Advanced Cell Diagnostics | 310043 |
| Nucleobond Xtra Midi Plus; DNA, RNA & protein purificatin | Machery-Nagel | 740412.5 |
| Opal 570 reagent pack | Akoya | FPI488001-KT |
| Positive control probe Mm-Ppib | Advanced Cell Diagnostics | 313911 |
| RNAScope multiplex Fluorescent V2 Assay Kit | Advanced Cell Diagnostics | 323100 |
| PVDF membrane | BioRad | 1620177 |
| Qubit RNA HS Assay Kit | Invitrogen | Q32855 |
| Seahorse XFe96 Flux pack | Agilent | 102416-100 |
| Target probe Mm-Dpt | Advanced Cell Diagnostics | 561511 |
| **Antibodies** |  |  |
| Anti Actin Fab- Rho | BioRad | 64422460 |
| Beta Actin Peroxidase Mouse mAb | Sigma | A3854 |
| Cytoglobin Rabbit pAb | Protein Tech | 13317-1-AP |
| Goat anti Mouse Alexaflour 488 | Invitrogen | A11029 |
| Goat anti Mouse-HPR | BioRad | 1705047 |
| Goat anti Rabbit Alexafluor 594 | Invitrogen | A32740 |
| Goat anti Rabbit StarBright Blue 700 | BioRad | 64247470 |
| Goat anti Rabbit-HPR | BioRad | 1705046 |
| Goat Serum | Vector labs | S-1000 |
| Mouse IgG isotype control | Vector labs | I-2000 |
| Peroxirecoxin-SO3 rabbit pAb | Abcam | ab16830 |
| Peroxiredoxin 1 Mouse mAb | Proteintech | 66820-1-Ig |
| Peroxiredoxin 1 Rabbit pAb | Proteintech | 20568-1-AP |
| Peroxiredoxin 2 Mouse mAb | Proteintech | 60202-1-Ig |
| Peroxiredoxin 2 Rabbit pAb | Proteintech | 10545-2-AP |
| Rabbit IgG isotype Control | Vector labs | I-1000 |

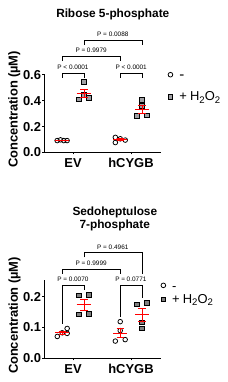

**A**

**B**

**Supplementary Figure 1. Effect of cytoglobin on the pentose monophosphate pathway.** Control HEK293 cells (EV = Empty Vector) and HEK293 cells expressing human cytoglobin (hCYGB) were treated with 150 µM hydrogen peroxide for 10 minutes. Concentrations of the pentose monophosphate pathway metabolites Ribose 5-phosphate (**A**) and Sedoheptulose 7-phosphate (**B**) were determined by mass spectrometry. All graphs show mean±SEM of four independent experimental replicates. Statistical analysis was performed using two-way ANOVA.

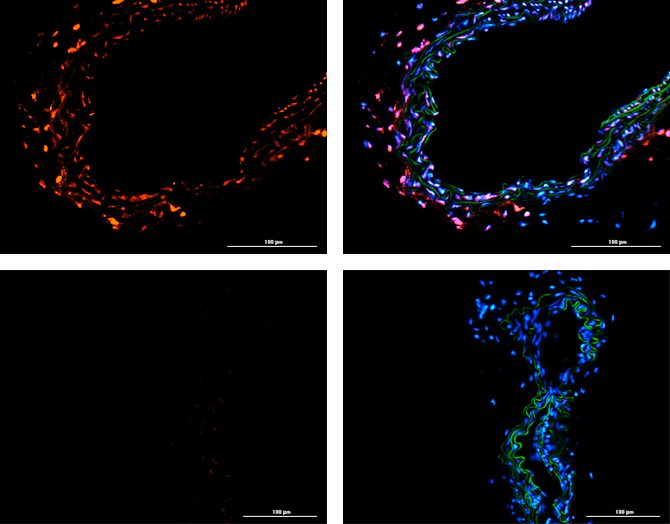

**C**

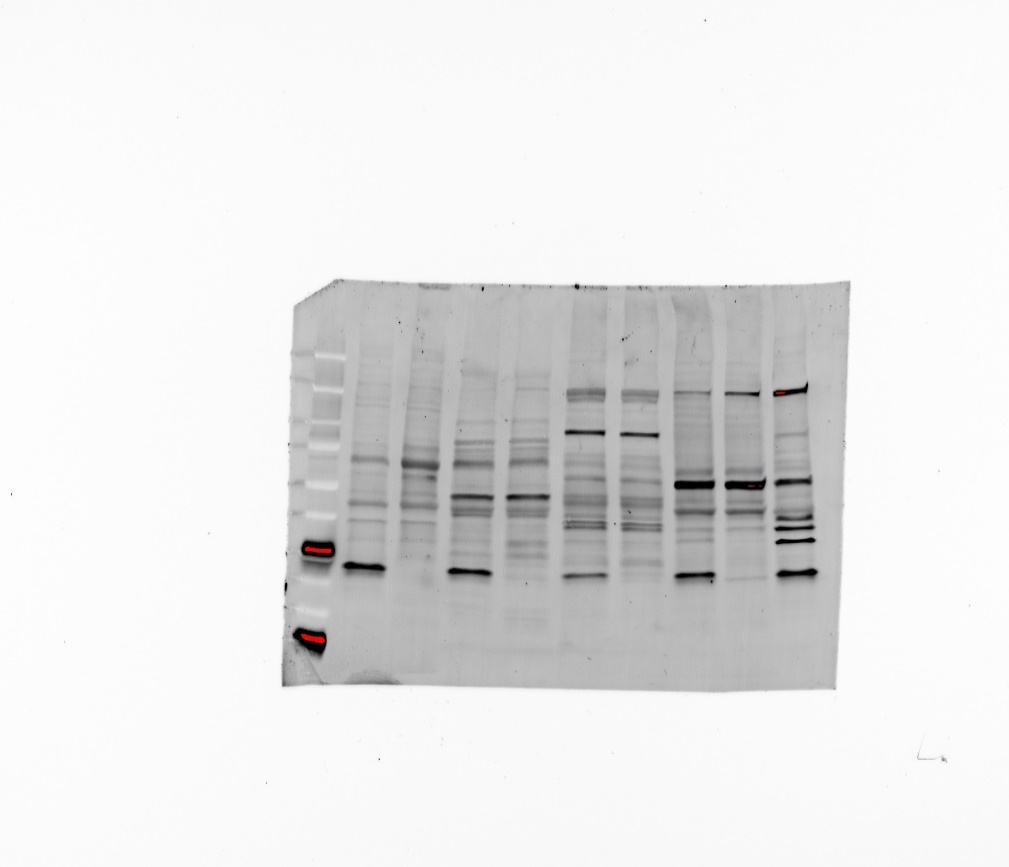

**IB: CYGB**

WT

KO

Spleen

Liver

Heart

Aorta

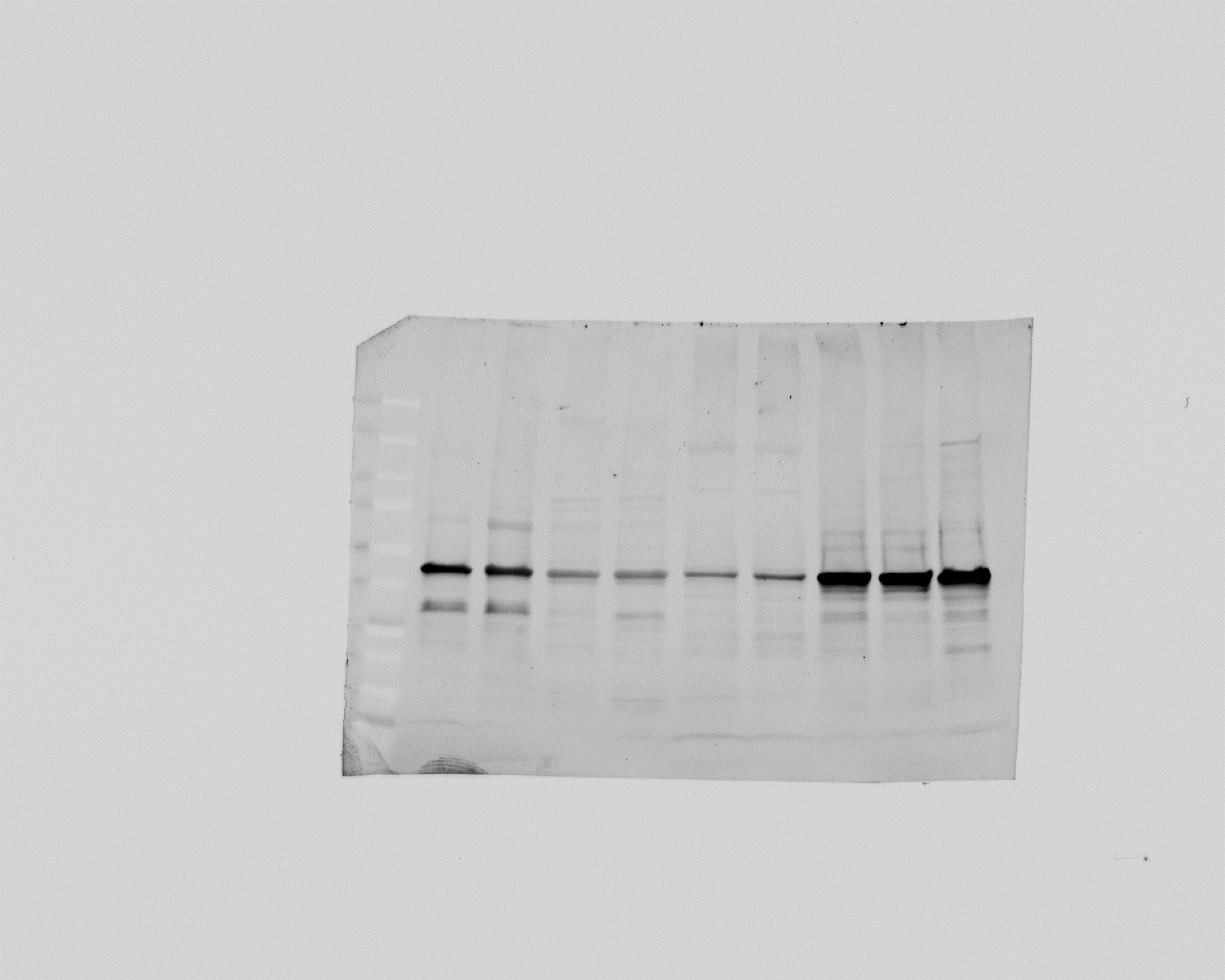

**IB: ACTB**

WT

KO

WT

KO

WT

KO

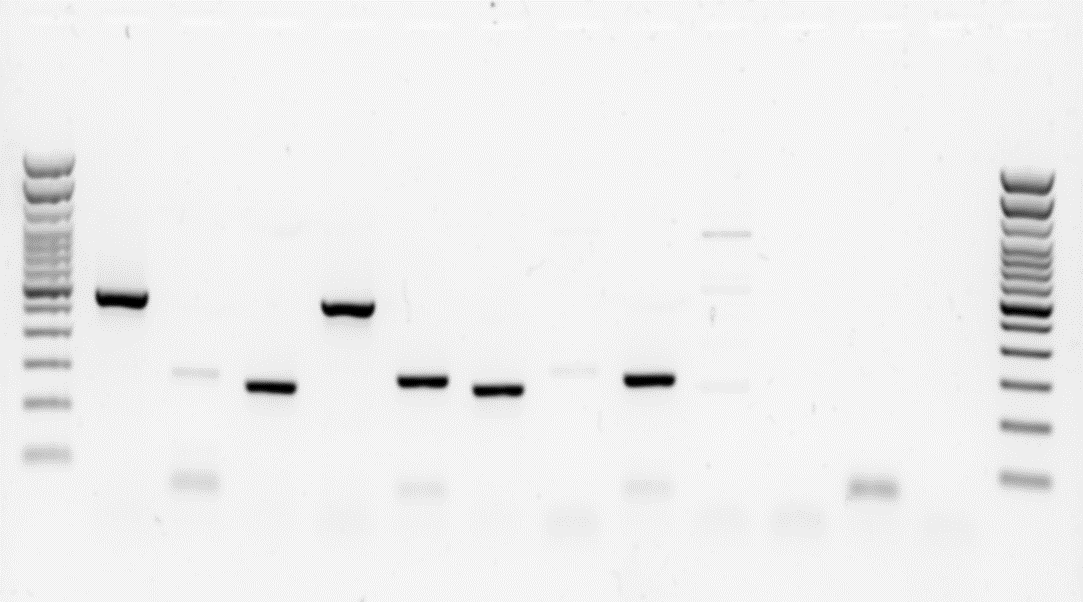

**CYGB KO**

**3**

**1**

**2**

**CYGB Het**

**F1155 (WT)**

**100**

**200**

**300**

**400**

**500**

**BP**

**Ladder**

**3**

**1**

**2**

**3**

**1**

**2**

**A**

**B**

Left Common Carotid Artery

CYGB WT

WT

Δ

FL

CYGB KO

**Supplementary Figure 2. Characterization of cytoglobin knockout mice. A,** Genotyping and recombination analysis of mouse tissues from global cytoglobin knockout (CYGB KO) wild-type (CYGB WT) and heterozygous (CYGB Het). Lane 1 contains excision reaction, lane 2 contains WT genotyping reaction, and lane 3 contains FL reaction. B, Representative Western blot for cytoglobin (CYGB) with β-actin (ACTB) as an internal reference in global cytoglobin knockout (KO) and wild-type (WT) mouse aorta, heart, liver, and spleen. C, Representative immunostaining for cytoglobin (red; blue is DAPI and green, autofluorescence) in tissue section from the left common carotid artery from global cytoglobin knockout (CYGB KO) wild-type (CYGB WT) mice.

**A**

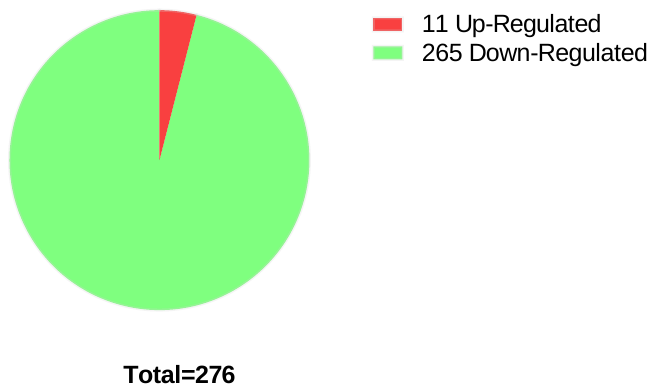

**Filter criteria:**

**Fold Change: < -1.25 or > 1.25**

**P-val: < 0.01**

**Cygb WT vs. Cygb KO**

**(Left common carotid artery; n=4 males)**

**B**

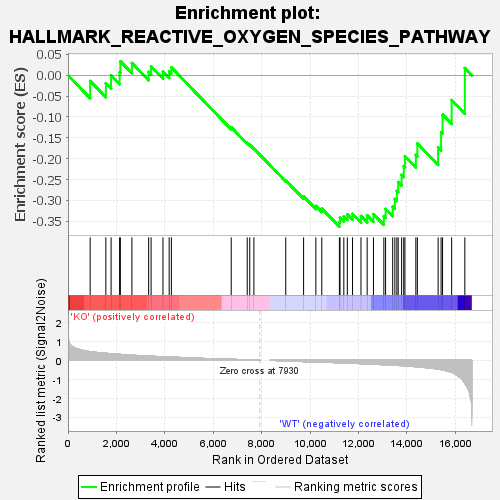

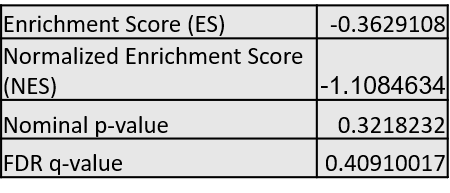

**C**

**Supplementary Figure 3. Bulk RNAseq in wild-type vs. cytoglobin mouse knockout left common carotid artery. A,** Venn diagram of the results of the RNA-seq analysis comparing normal left common carotid arteries of CYGB WT and CYGB KO. Differentially expressed genes (DEGs) were selected with the criteria of fold-change < −1.25 or > 1.25, p-value < 0.01. **B,** GSEA enrichment plot for reactive oxygen species pathway using RNA-seq data comparing normal left common carotid arteries of CYGB WT and CYGB KO. **C,** GSEA enrichment results for reactive oxygen species pathway using RNA-seq data comparing normal left common carotid arteries of CYGB WT and CYGB KO.

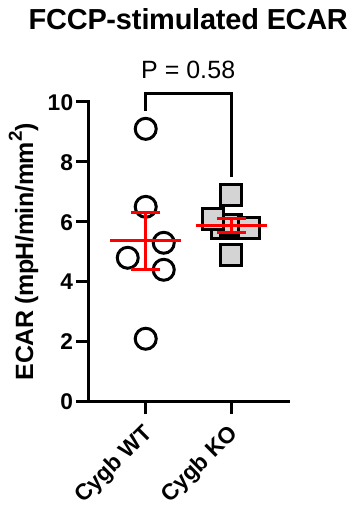

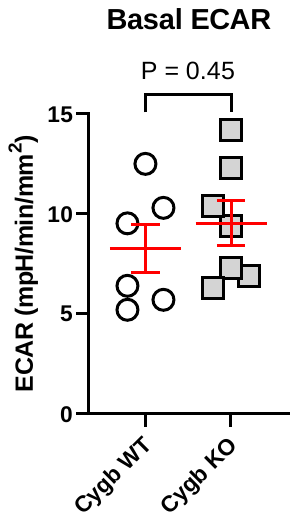

**A**

**B**

**Supplementary Figure 4. A,** Basal ECAR obtained from isolated common carotid arteries from Cygb WT and Cygb KO mice**. B,** FCCP-stimulated ECAR from isolated common carotid arteries from Cygb WT and Cygb KO mice. Each data point indicates one mouse. Statistical analysis was performed using unpaired Student’s t-test
